# Cell-to-cell heterogeneity drives host-virus coexistence in a bloom-forming alga

**DOI:** 10.1101/2023.10.03.560477

**Authors:** Nir Joffe, Constanze Kuhlisch, Guy Schleyer, Nadia Samira Ahlers, Adva Shemi, Assaf Vardi

## Abstract

Algal blooms drive global biogeochemical cycles of key nutrients in the oceans and serve as hotspots for biological interactions. The massive spring blooms of the cosmopolitan coccolithophore *Emiliania huxleyi (E. huxleyi)* are often infected by the lytic *Emiliania huxleyi* specific virus (EhV) which is a major mortality agent triggering bloom demise. Nonetheless, the multi-annual “boom and bust” pattern of *E. huxleyi* suggests that mechanisms of coexistence are essential for these host-virus dynamics. To investigate host-virus coexistence, we developed a new model system from an *E. huxleyi* culture which recovered from viral infection. The recovered population coexists with the virus, as host cells continue to grow in parallel to viral production. By applying a single-molecule fluorescence *in situ* hybridization (smFISH) approach to quantify the fraction of infected cells and assessing infection-specific lipid biomarkers, we identified a small subpopulation (5-7% of cells) that was infected and produced new virions, whereas the majority of the host population could resist infection. To further assess population heterogeneity, we generated monoclonal strain collections using single-cell sorting and subsequently phenotyped their susceptibility to EhV infection. This unraveled a substantial cell-to-cell heterogeneity across a continuum of susceptibility to resistance, suggesting that infection outcomes may vary depending on the individual cell. These results add a new dimension to our understanding of the complexity of host-virus interactions that are commonly assessed in bulk and described by binary definitions of resistance or susceptibility. We propose that phenotypic heterogeneity drives *E. huxleyi*-EhV coexistence and may potentially provide the coexisting strain an ecological advantage by killing competing susceptible strains.

## Introduction

Algal blooms are ephemeral events of massive cell proliferation, serving as ecological hotspots of primary production and microbial interactions in the ocean (1). Marine viruses are a major factor controlling algal blooms. By infecting their hosts, viruses reduce the host population size, reshape their composition, and influence nutrient and organic carbon cycling (2). Infection strategies of viruses vary from lytic infection to non-lethal infections such as lysogeny and chronic infection. Due to host replication dependency, host extinction is thereby an undesirable outcome. Non-lethal infections can ensure host survival as the virus can persist intracellularly without hindering cell division. In contrast, lytic viruses lead to rapid host cell death and virion release, preventing host-virus coexistence at the single-cell level (3–5). This raises a key ecological question of how hosts and lytic viruses coexist and how algal blooms reoccur throughout the years. Although several studies showed algal-virus coexistence (6–9), we still lack understanding of this process, especially its dynamics at a single-cell resolution and the effect on population heterogeneity.

The cosmopolitan coccolithophore *Emiliania huxleyi* and its specific virus, the *E. huxleyi* virus (EhV), is an important host-virus model system with significant ecological impact. *E. huxleyi* forms vast annual spring blooms in temperate regions that can cover thousands of square kilometers (10). These large-scale blooms are often infected by EhV that acts as a dominant mortality agent causing bloom termination (11–13). EhV is a large (ca. 180 nm in diameter), double-stranded DNA virus with a lytic life cycle and high burst size (14). Despite its virulent nature, EhV does not lead to *E. huxleyi* extinction, as evidenced by the multi-annual cycle of bloom and demise. This dilemma becomes more complex when the virus is specialized on a bloom-forming alga, where host availability is limited to the high cell densities during ephemeral bloom events. This specialized strategy of EhV is very challenging in the long periods lasting many months in between blooms, especially with regard to the half-life of virions in the aquatic ecosystem leading to a daily decrease in their infectivity (15–18). It has been shown that *E. huxleyi* strains vary in their susceptibility to different EhV isolates and that most *E. huxleyi* strains are resistant to some if not all, EhV isolates (14, 19). This prominent occurrence of virus resistance among *E. huxleyi* strains suggests a complex host-virus interaction that prevents the eradication of the virus over evolutionary time scales. Although *E. huxleyi* and its lytic virus coexisted for thousands of years in the natural environment (20), the ecological and biological processes facilitating their coexistence are underexplored (21). Previous studies suggest that *E. huxleyi*-EhV dynamics follow the continuous arms race model, whereby both players genetically adapt to overcome one another. A central manifestation for such coevolution is the large arsenal of auxiliary metabolic genes (AMGs) that are encoded by EhV and transcribed during infection. AMGs rewire the host cell metabolism and thereby facilitate the production of essential building blocks for new viral progeny. For example, EhV encodes an almost complete sphingolipid pathway leading to the biosynthesis of virus-derived glycosphingolipids (vGSLs) during viral infection (22–24). The viral genes are phylogenetically close to the corresponding host genes, indicating horizontal gene transfer as part of the evolutionary arms race between *E. huxleyi* and EhV (25). In addition, *E. huxleyi* may evade viral attack by morphological and life-cycle changes within a small subpopulation of cells that promote resistance (26, 27). Nevertheless, infection experiments in the lab typically end with one winner, either EhV or a resistant *E. huxleyi* strain (26–31). We thus lack an experimental model system to study host-virus coexistence.

We aimed to investigate the occurrence of host-virus coexistence in the *E. huxleyi*-EhV model system, its effect on infection dynamics, and its link to host phenotypic heterogeneity. We examined the interaction of the susceptible *E. huxleyi* strain CCMP 2090 (hereinafter, *E. huxleyi* 2090), and the lytic virus strain EhV-201 in a lab-based experiment. Specifically, we characterized a resistant *E. huxleyi* culture that recovered from viral infection, named *E. huxleyi* 2090-Rec, which maintained a parallel proliferation of both the host and virus. We isolated 74 clonal cultures that were derived from single-cell isolations of *E. huxleyi* 2090-Rec and phenotyped their susceptibility to viral infection. In contrast to an expected binary resistance or susceptibility in these clones, we revealed a wide spectrum of resistance levels across these single-cell isolates. These results highlight the cell-to-cell heterogeneity within host populations and provide a new perspective on the binary definition of resistance and susceptibility at the population level. We propose that the phenotypic plasticity of *E. huxleyi* is the driving force for establishing coexistence with its lytic virus and suggest how the multi-annual dynamic of *E. huxleyi* bloom and EhV bloom termination is sustained without host or virus extinction.

## Results

### Development of a model system for studying host-virus coexistence

To test whether the microalga *E. huxleyi* and its specific virus EhV can establish coexisting populations under controlled culture conditions, the susceptible *E. huxleyi* 2090 was infected with the lytic viral strain EhV-201 (Fig. 1A). The infection dynamics were monitored for 40 days by counting host and virus abundances using flow cytometry (Fig. 1B). At 2-11 days post-inoculation (dpi), the host population showed a virus-induced decline of 99.9% of the cells (Fig. 1B). The population was not lysed completely and remained at a minimal cell density of ∼10^3^ cells mL^-1^ for six days. At 20 dpi, the recovery phase started, characterized by increasing cell abundances that exceeded the initial host population density at 37 dpi. The host population decline in the first 3 dpi was coupled with high viral production (Fig. 1B) and up to 80% cell death at 6 dpi (Supplementary Fig. S1). The recovered host population grew in the presence of substantial viral load (6×10^7^-1×10^8^ viral particles mL^-1^), suggesting the development of phenotypic properties that differ from the susceptible ancestor strain and provide resistance to viral infection. To examine virus resistance in the recovered host population, the cultures were diluted weekly into fresh medium, reaching a dilution of at least 10^8^, to minimize the virions of the viral lysate (Fig. 1A). Notably, virus propagation (>1×10^8^ viral particles mL^-1^) was detected in parallel to algal cell growth in these cultures, reaching a maximum of 4×10^6^ cells mL^-1^ in the stationary phase (Fig. 1C). The emergence of a recovered resistant cell population, named *E. huxleyi* 2090-Rec (2090-Rec), together with the proliferation of a co-occurring virus, named EhV-Rec, suggests the development of a stable coexistence between *E. huxleyi* and EhV. We then examined whether EhV-Rec exhibits different infectivity towards *E. huxleyi* strains as compared to the ancestor EhV-201. The infectivity of EhV-Rec towards the susceptible *E. huxleyi* strains 2090, CCMP374 (hereinafter, 374), and RCC1216 (hereinafter,1216), as well as the resistant strain CCMP379 (hereinafter, 379), was similar to EhV-201 (Supplementary Fig. S2). Thus, EhV-Rec exhibited a similar host range and infection dynamics as EhV-201, suggesting that no major phenotypic differences exist between the virions at infection phase compared to the coexistence state. Furthermore, virus resistance in the recovered *E. huxleyi* population 2090-Rec was tested against the virus strains EhV-201, EhV-Rec, EhV-86, EhV-163, and EhV-M1, which differ in their host range and infection dynamics with the ancestral *E. huxleyi* strain 2090 (Fig. 1D-E). In contrast, none of these viruses affected the proliferation of 2090-Rec cells (Fig. 1D, *p*-value =0.0931) or led to viral production (Supplementary Fig. S3A) (*p*-value =0.1207). When inoculated with EhV-201, EhV-Rec, or EhV-86, cultures of the ancestral *E. huxleyi* strain 2090 declined (Fig. 1E), accompanied by an increase in virus abundance (Supplementary Fig. S3B) as previously shown (32–34). Taken together, we observed similarities between the infection dynamics governed by the coexisting virus EhV-Rec compared to its ancestor strain EhV-201, and dissimilarities in the susceptibility of the coexisting host population *E. huxleyi* 2090-Rec compared to its ancestor strain *E. huxleyi* 2090. Therefore, we hypothesize that the generation of a host-virus coexistence state is driven by changes in the host population rather than in the virions themselves.

**Figure 1.**
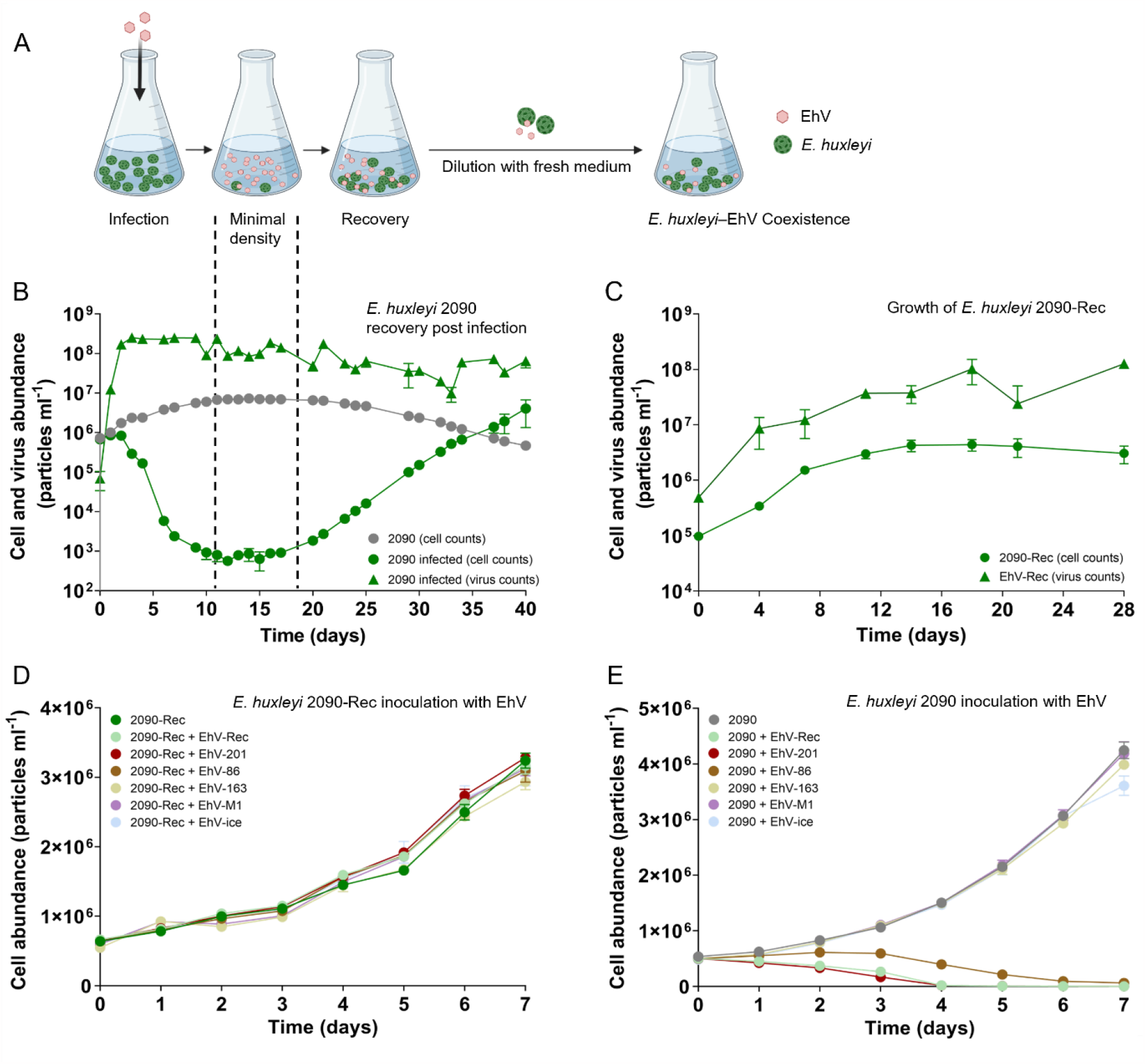
Coexistence of lytic viruses with a population of *E. huxleyi* cells recovered from viral infection. (A) Schematic representation of the experimental system. Upon viral infection, *E. huxleyi* populations decline to a small number of cells that survive and subsequently form a new population. This recovered resistant population proliferates in the presence of lytic viruses and produces virions over many generations. (B) Infection of *E. huxleyi* 2090 with EhV-201 led to the recovery of the resistant population *E. huxleyi* 2090-Rec in the presence of lytic viruses. (C) Concomitant proliferation of host cells and virions in cultures of the recovered coexisting *E. huxleyi* 2090-Rec following continuous sub-cultivation in fresh growth medium. (D) Growth of *E. huxleyi* 2090-Rec cells is not affected by inoculation with different EhV strains. (*E*) Growth of ancestral *E. huxleyi* 2090 cells upon inoculation with different EhV strains. Values are presented as mean ± SD (n = 3).

### Coexisting recovered cell populations are heterogenic and composed of subpopulations

*E. huxleyi*-EhV interactions have been assessed mainly by bulk measurements, assuming phenotypic homogeneity of the cells within a population. The observed coexistence of a resistant host population and a lytic virus suggests the co-occurrence of phenotypically diverse cells within the host population. We thus sought to assess the phenotypic heterogeneity within the *E. huxleyi* 2090-Rec population in comparison to a population of the ancestral strain *E. huxleyi* 2090 using established molecular and metabolic markers for viral infection in *E. huxleyi* (23, 35). We first applied lipid biomarkers (GSLs) that can inform about the phenotypic cell states of *E. huxleyi*, namely uninfected cells (host-derived GSLs, hGSLs), susceptible cells (sialic acid GSLs, sGSLs), and virus-infected cells (vGSLs) (23, 35). Exponentially growing cultures of *E. huxleyi* 2090-Rec were analyzed using a liquid chromatography-high-resolution mass spectrometry (LC-HRMS)-based lipidomics approach and compared to infected and uninfected cultures of the ancestral strain *E. huxleyi* 2090. Cell cultures of the coexisting *E. huxleyi* 2090-Rec were comprised of hGSLs, sGSLs and vGSLs similar to virus-infected *E. huxleyi* 2090, while uninfected cultures of the ancestral strain comprised only hGSLs and sGSLs (Fig. 2A; Supplementary Fig. S4). The detection of vGSL in *E. huxleyi* 2090-Rec cultures indicates the presence of a subpopulation of infected cells, and due to the stable growth of the *E. huxleyi* 2090-Rec cultures, we assume the population is composed of another subpopulation of uninfected resistant cells.

**Figure 2.**
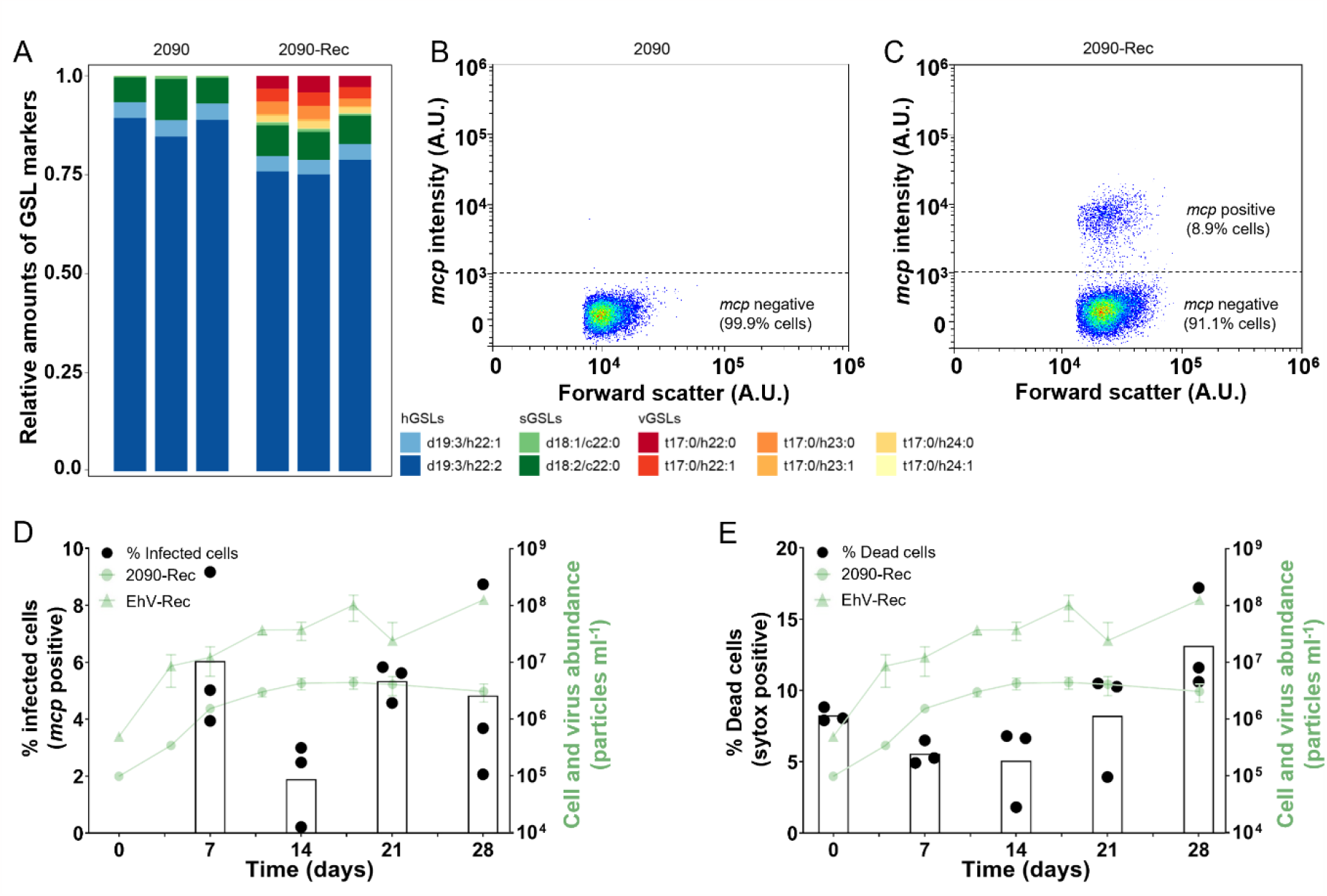
The recovered coexisting *E. huxleyi* 2090-Rec population comprises a subpopulation of virus-infected cells undergoing cell lysis. (A) Relative amounts of GSL markers (hGSLs, sGSLs and vGSLs) in *E. huxleyi* 2090 and 2090-Rec. (B) Single cell expression of the viral *mcp* gene using smFISH in the recovered coexisting culture *E. huxleyi* 2090-Rec during late exponential growth at day 7. Using a threshold intensity of 10^3^ A.U. for the *mcp* probe, 8.9% cells expressed the viral *mcp* gene (n = 17,612). (C) Single cells from an uninfected 2090 culture show no expression of the viral *mcp* gene (n = 13,942). Boxplots depict the fraction of (D) infected cells (*mcp* positive cells, as quantified by smFISH) and (E) dead cells (Sytox positive) in cultures of *E. huxleyi* 2090-Rec throughout growth (left y-axis). Green lines depict counts of cells 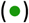 and viruses 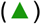 (right y-axis), as in Fig. 1C. Values are presented as mean ± SD (n = 3).

To further quantify these subpopulations in *E. huxleyi* 2090-Rec, we used single-molecule fluorescence *in situ* hybridization (smFISH), which enables quantification of viral transcripts at a single-cell resolution and informs about the fraction of infected cells in a population (4). Flow cytometry was used to detect and enumerate cells with a positive fluorescence signal using probes that target mRNA transcripts of the viral major capsid protein (*mcp)* gene (Fig. 2A and B). In *E. huxleyi* 2090-Rec, the fraction of cells that were actively infected (*mcp* positive) was on average 2-6% throughout exponential and stationary growth (Fig. 2D), while the fraction of dead cells was 5-13% (Fig. 2E). In comparison, uninfected cultures of *E. huxleyi* 2090 showed only *mcp* negative cells (Fig. 2B, Supplementary Fig. S5) and 2-4% of dead cells (Supplementary Fig. S1). Intriguingly, these results demonstrate that the coexisting *E. huxleyi* 2090-Rec is a heterogeneous population comprising at least two subpopulations with contrasting phenotypes: one being resistant and one infected. While bulk cell abundance measurements suggest that the *E. huxleyi* 2090-Rec population is overall resistant, the presence of vGSLs, *mcp* positive cells and elevated levels of dead cells indicate that a small fraction of cells in this resistant recovered population is virus-susceptible and undergoing lytic viral infection.

### Mapping population heterogeneity in resistance to viral infection at a single-cell level

To explore the heterogeneity in virus resistance within the recovered coexisting *E. huxleyi* population compared to the ancestor strain, a single-cell sorting approach was applied. Monoculture collections were generated by sorting single cells originating from either coexisting or susceptible cultures into well plates (Fig. 3A). In total, 485 single cells were isolated from *E. huxleyi* 2090-Rec and 388 single cells from the ancestor strain *E. huxleyi* 2090, of which 123 and 74, respectively, formed viable growing monocultures (Supplementary Table S1). This monoculture collections represent, to some extent, the cell heterogeneity of the population from which they were derived. Therefore, observing differences between individual monocultures reflects the cell-to-cell phenotypic heterogeneity within the original cell population. To test for the presence of viral particles and thus of possible host-virus coexistence, the abundance of extracellular EhV in all monocultures was measured by quantitative PCR (qPCR) using primers for the EhV *mcp* gene (Supplementary Fig. S6). Notably, none of the monocultures that were derived from *E. huxleyi* 2090-Rec produced viruses. Subsequently, all monocultures were phenotyped for virus resistance by monitoring cell abundances (based on chlorophyll fluorescence) and measuring the virus production six days following the addition of EhV-201 (Fig. 3B and C).

**Figure 3.**
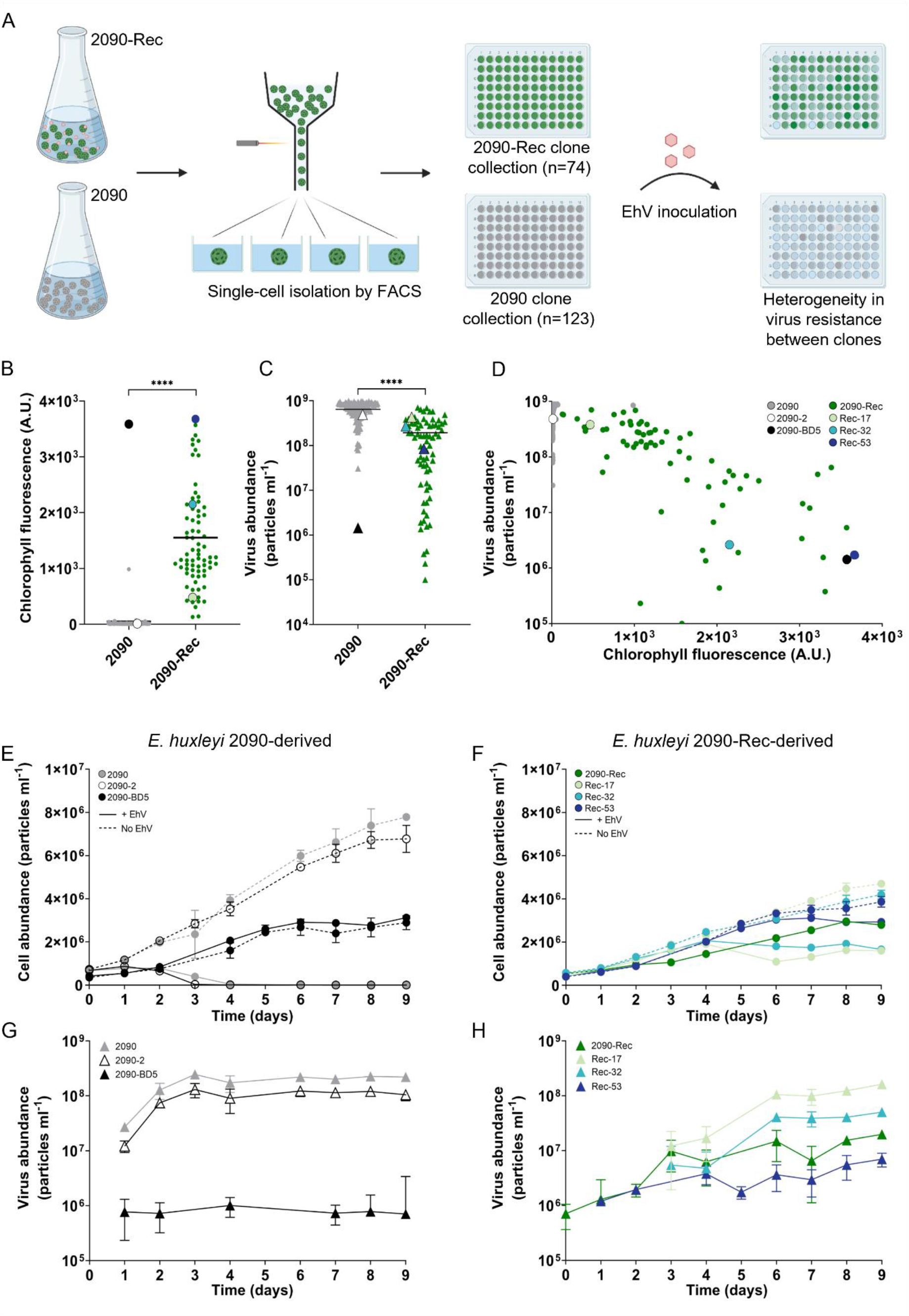
Cell-to-cell heterogeneity in resistance to viral infection within coexisting *E. huxleyi* 2090-Rec. (A) Experimental procedure of the single-cell isolation and phenotyping for viral resistance. Single cells were sorted from susceptible strain *E. huxleyi* 2090 and coexisting culture *E. huxleyi* 2090-Rec, and the derived monocultures were inoculated with the virus EhV-201. The resistance phenotype of each monoculture was characterized at day 6 of the experiment. (B) Chlorophyll fluorescence of monocultures derived from *E. huxleyi* 2090 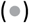 and 2090-Rec 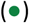 is used as a proxy for culture growth and survival during viral infection. (C) Viral particle abundance in monocultures derived from *E. huxleyi* 2090 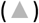 and 2090-Rec 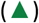. (D) Virus abundance as a function of chlorophyll fluorescence in monocultures derived from *E. huxleyi* 2090 and 2090-Rec. Monocultures with different levels of virus resistance were selected for further investigation, namely monocultures 2090-2 (?) and 2090-BD5 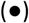 derived from *E. huxleyi* 2090, and monocultures Rec-17 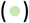, Rec-32 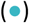 and Rec-53 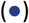 derived from *E. huxleyi* 2090-Rec. (E) Cell abundance of virus-inoculated (full lines) and non-inoculated (dashed lines) *E. huxleyi* 2090 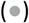 cultures and *E. huxleyi* 2090-derived monocultures. (F) Cell abundance of *E. huxleyi* 2090-Rec 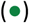 and virus-inoculated (full lines) and non-inoculated (dashed lines) *E. huxleyi* 2090-Rec-derived monocultures. (G) Virus production in virus-inoculated *E. huxleyi* 2090 and 2090-derived monocultures. (H) Virus production in *E. huxleyi* 2090-Rec and virus-inoculated *E. huxleyi* 2090-Rec-derived monocultures. Values in E-H are presented as mean ± SD (n = 3).

The mean chlorophyll fluorescence of *E. huxleyi* 2090-Rec-derived monocultures was two orders of magnitude higher than that of *E. huxleyi* 2090-derived monocultures (*p*-value <0.0001, 1.5×10^3^ vs. 10 A.U., Fig. 3B), reflecting their resistance to viral infection. Accordingly, the mean virus production of monocultures derived from *E. huxleyi* 2090 at 6 dpi was ∼4-fold higher than that of the *E. huxleyi* 2090-Rec-derived monocultures (*p*-value <0.0001, 6.5×10^8^ vs. 1.5×10^8^ viral particles mL^-1^, Fig. 3C). Furthermore, the heterogeneity in virus production was larger in monocultures derived from *E. huxleyi* 2090-Rec compared to monocultures derived from *E. huxleyi* 2090, with a coefficient of variance 99.7% vs. 39.1%. Similarly, *E. huxleyi* 2090-Rec-derived monocultures showed a high diversity in chlorophyll fluorescence, while the majority of *E. huxleyi* 2090-derived monocultures were dead, with only two viable monocultures that showed a chlorophyll fluorescence of >10^2^ A.U. These results indicate that *E. huxleyi* 2090-Rec comprises a highly heterogeneous population of cells that vary in their susceptibility while *E. huxleyi* 2090 is comprised of a mostly homogeneous population of susceptible cells. The concomitant occurrence of resistant monocultures having high chlorophyll fluorescence and low viral production, and monocultures with lower virus resistance, having low chlorophyll fluorescence and high viral production, provide evidence for phenotypic heterogeneity (Fig. 3D). The chlorophyll fluorescence and virus production of *E. huxleyi* 2090-Rec monocultures were negatively correlated (r = -0.784, *p*-value <0.0001). These results suggest that the coexisting *E. huxleyi* 2090-Rec population is composed of cells with phenotypes across a continuum of susceptibility rather than in a binary definition of being either susceptible or resistant. Noteworthy, the presence of two monocultures from the susceptible ancestor *E. huxleyi* 2090, which were resistant to viral infection, highlights that viral resistance can also emerge from susceptible populations in the absence of viral pressure leading to subpopulations of resistant cells.

To further characterize different resistant cells that co-occur in the *E. huxleyi* 2090-Rec population, we selected monocultures with distinct resistance phenotypes. The monocultures Rec-17, Rec-32, and Rec-53 were selected as they span the resistance-susceptibility-continuum, with Rec-17 being most susceptible and Rec-53 being most resistant (Fig. 3B-D). In addition, two monocultures derived from the susceptible strain *E. huxleyi* 2090 were selected: the monoculture 2090-BD5, as an emerged resistant monoculture, and 2090-2, as a representative susceptible monoculture (Fig. 3B-D). The selected monocultures were picked from the 96-well plates, grown in larger culture volumes, and inoculated with EhV-Rec. The infection dynamics were monitored at high temporal resolution by measuring the abundance of cells and virus particles (Fig. 3E-H). Virus-inoculated cultures of monoculture 2090-2 demonstrated growth arrest at 2 dpi and most of the population lysed within 7 dpi, similar to the ancestor *E. huxleyi* 2090 (Fig. 3E). This was coupled with high virus production in both *E. huxleyi* 2090 and 2090-2 starting from 2 dpi (∼1×10^8^ viral particles mL^-1^, Fig. 3G). In contrast, virus-inoculated cultures of 2090-Rec, Rec-17, Rec-32, Rec-53, and 2090-BD5, did not exhibit an intense population decline in parallel to the production of viruses (Fig. 3E-H). Noteworthy, monocultures 2090-BD5, Rec-53, Rec-32, and Rec-17 displayed growth together with virus production, suggesting the existence of a fraction of susceptible cells within their population. We confirmed their phenotypic heterogeneity through single-cell resistance screening, thus demonstrating the ability of individual cells to generate diverse populations (Supplementary Fig. S7). The maximum viral production (MVP, Supplementary Fig. S8), maximum carrying capacity (CC, Supplementary Fig. S9), and growth rate (µ, Supplementary Fig. S10) of all cultures were computed to characterize the *E. huxleyi*-EhV infection dynamics and differentiate the levels of virus resistance of these strains. The MVP varied across the virus-containing monocultures, Rec-17 and 2090-2 had a similar MPV to that of the ancestor *E. huxleyi* 2090 (1.6×10^8^ viral particles mL^-1^), while the other monocultures had lower MPV values compared to *E. huxleyi* 2090: Rec-32 was ∼70% lower (0.5×10^8^ viral particles mL^-1^), Rec-53 was ∼96% lower (7×10^6^ viral particles mL^-1^), 2090-BD5 was 99.6% lower (1×10^6^ viral particles mL^-1^), and also the recovered coexisting *E. huxleyi* 2090-Rec was ∼88% lower than *E. huxleyi* 2090 (2×10^7^ viral particles mL^-1^, Supplementary Fig. S8). In the presence of viruses, the monocultures Rec-53, 2090-BD5 and 2090-Rec demonstrated high CCs (2.9×10^6^, 3.1×10^6^, 3.3×10^6^ cells mL^-1^, respectively), Rec-17 and Rec-32 displayed lower CCs (2×10^6^ cells mL^-1^), while the susceptible *E. huxleyi* 2090 and 2090-2 showed a low CC (<1×10^6^ cells mL^-1^), suggesting that a high CC in the presence of viruses illustrates a high resistance level (Supplementary Fig. S9A). Furthermore, the non-inoculated monocultures Rec-17, Rec-32, Rec-53, and 2090-BD5 displayed a lower CC and µ than non-inoculated *E. huxleyi* 2090 and 2090-2, highlighting low CC and growth rates as a cost of resistance (Fig. 3E, G and Supplementary Fig. S9B and S10A). Taken together, by assessing diverse phenotypes originating from the same population, we propose that *E. huxleyi* can generate cells with different resistance levels. Characterizing virus resistance at single-cell resolution unveiled substantial cell-to-cell heterogeneity within the *E. huxleyi* 2090-Rec population. This heterogeneity allows the host to remain resistant to lytic viral infection at the population level while coexisting with high phenotypic heterogeneity, including susceptible cells. This coexistence enables constant virus production while maintaining resistance at the population level.

### Benefits of host-virus coexistence during intraspecies competition

To assess possible ecological consequences of *E. huxleyi*-EhV coexistence in mixed natural bloom populations, we conducted a series of co-cultivation assays simulating intraspecies competition against other host strains that differ in their virus resistance. The coexisting *E. huxleyi* 2090-Rec was co-cultured with two strains that are susceptible to viral infection (*E. huxleyi* 2090 and 374), one strain that is fully resistant (*E. huxleyi* 379), and *E. huxleyi* 2090-BD5 that is overall resistant while showing minor viral production after EhV inoculation. The cultures were separated by a 1 µm pore-size filter, allowing the exchange of viruses, nutrients and metabolites, but not of algal cells (Fig. 4A).

**Figure 4.**
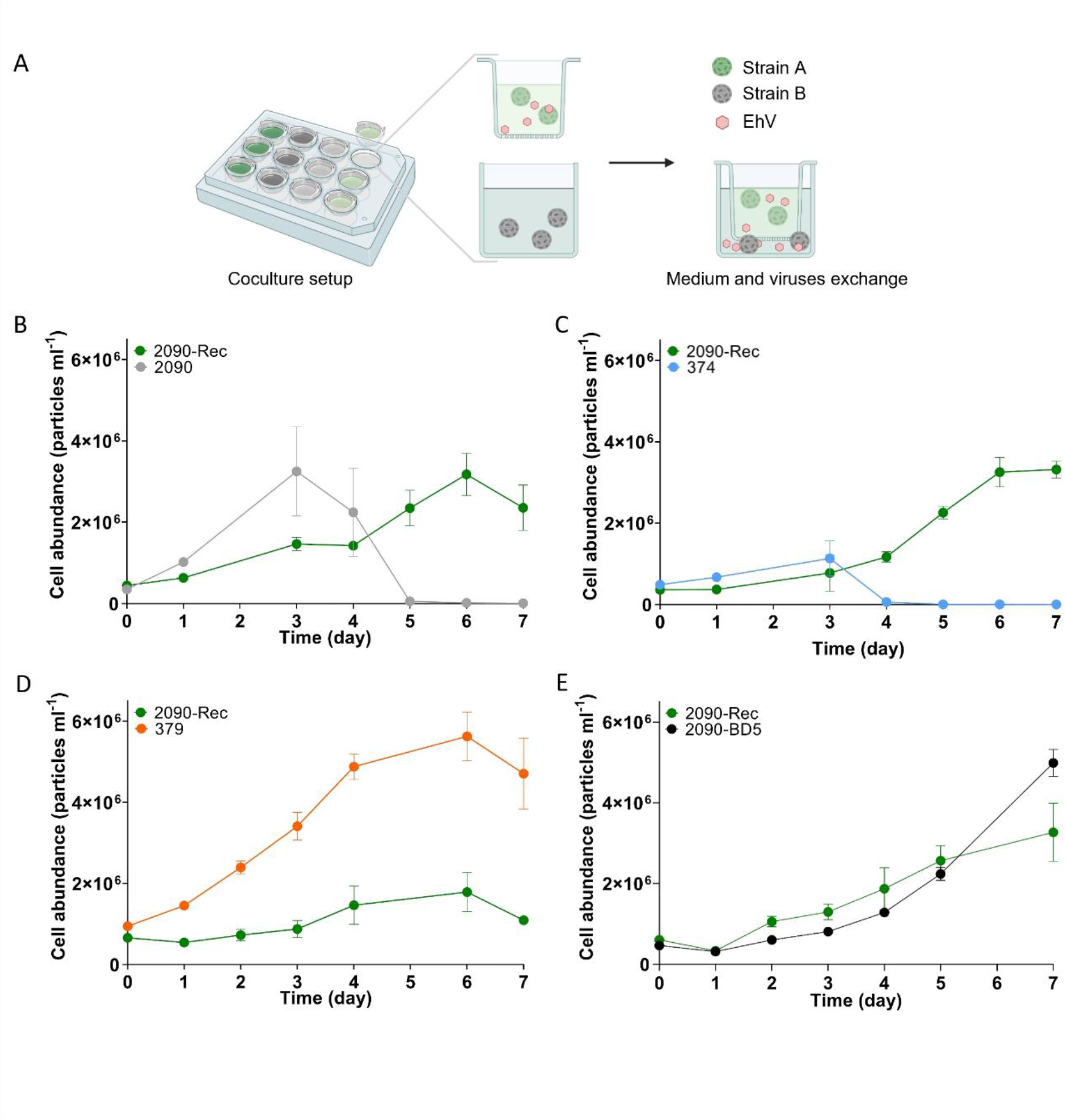
Viral production in host-virus coexistence can benefit against competing susceptible host strains. (A) Schematic representation of the co-cultivation setups. Couples of *E. huxleyi* strains were co-cultured in 24-well plates separated by a 1 µm membrane, allowing the exchange of nutrients, metabolites, and viruses, but preventing the mixing of algal cells. Cell abundances of the coexisting *E. huxleyi* 2090-Rec during co-cultivation with (B) *E. huxleyi* 2090, (C) 374, (D) 2090-BD5, and (E) 379. Strain 2090-Rec was grown in the insert and the competing strains within the surrounding wells. Values in B-E are presented as mean ± SD (n = 3).

During co-cultivation of *E. huxleyi* 2090-Rec with its susceptible ancestor *E. huxleyi* 2090, *E. huxleyi* 2090 outgrew the coexisting population in the first three days, reflecting a higher growth rate and a possible trade-off for resistance. However, in the subsequent days, the *E. huxleyi* 2090 population rapidly declined due to infection by the viruses (EhV-Rec) that were released from *E. huxleyi* 2090-Rec (Fig. 4B). These results emphasize how the slow-growing *E. huxleyi* 2090-Rec outcompetes the fast-growing susceptible *E. huxleyi* 2090 by continuous release of infectious virions. The viruses that are produced by the coexisting population served as a weapon against the susceptible strain leading to its decline and more viral progeny (Supplementary Fig. S11A). Similar results were observed by co-culturing *E. huxleyi* 2090-Rec with the susceptible strain *E. huxleyi* 374 (Fig. 4C; Supplementary Fig. S11B). On the other hand, when co-cultured with the resistant strain *E. huxleyi* 379, *E. huxleyi* 2090-Rec was outgrown (Fig. 4D). *E. huxleyi* 379 reached a substantially higher cell density compared to *E. huxleyi* 2090-Rec (5.6x10^6^ cells mL^-1^ and 1.8x10^6^ cells mL^-1^, respectively), suggesting that heterogeneity and host-virus coexistence is not a favorable competition strategy against a homogenous resistant algal population. Nevertheless, the abundances of resistant strains are typically extremely low in *E. huxleyi* blooms (36).

When *E. huxleyi* 2090-Rec was co-cultured with *E. huxleyi* 2090-BD5, the coexisting *E. huxleyi* 2090-Rec grew to similar cell densities as *E. huxleyi* 2090-BD5 (Fig. 4E). This result indicates that there is no fitness advantage for either strain when co-occurring in the same population and that EhV-Rec produced by *E. huxleyi* 2090-Rec (Supplementary Fig. S11C) had no significant effect on the growth of *E. huxleyi* 2090-BD5 (*p*-value=0.6986). Noteworthy, in the absence of viruses, the susceptible strain *E. huxleyi* 2090 outcompeted the resistant *E. huxleyi* 2090-BD5 (Supplementary Fig. S12F), highlighting the cost of resistance in a virus-free environment. This cost of resistance may explain why so few resistant cells were isolated from *E. huxleyi* 2090.

Taken together, the heterogeneous *E. huxleyi* 2090-Rec population represents a mutualistic host-virus interaction. The coexistence with a lytic virus is advantageous for *E. huxleyi* during intraspecies competition via proliferating viruses produced by a minor subpopulation whereas the rest of the population is resistant. Interestingly, we found host-virus coexistence to emerge in several other *E. huxleyi*-EhV interactions including well-studied strains as well as recent isolates from *E. huxleyi* mesocosm blooms near Bergen in Norway (Supplementary Fig. S13), suggesting host-virus coexistence to be a common and widespread phenomenon.

## Discussion

Host extinction is an undesirable outcome for viruses due to their dependency on replication within host cells. Different strategies can ensure the coexistence of a virus with its host, including lysogeny, chronic infection, and mutualistic interaction, where virus infection promotes host competitiveness (37, 38). However, mechanisms that allow a host to coexist with its lytic virus are still underexplored. This dilemma becomes even more complex when the virus is specialized on bloom-forming algae whose availability is limited to the high cell densities during ephemeral bloom events.

### Heterogeneity within the host population is a driving force for host-virus coexistence

When the entire *E. huxleyi* population shares uniform susceptibility to the virus, an encounter with EhV should lead to the lysis of all cells and host extinction. Conversely, if all cells are uniformly resistant, there will be no virus progeny leading to virion decay and virus extinction. Host populations exhibiting these binary characteristics will not be able to maintain coexistence with a lytic virus. The observed phenotypic heterogeneity in *E. huxleyi* led us to hypothesize that host-virus coexistence is driven by the continuous formation of heterogeneous cells characterized by different resistance levels. The emergence of a more resistant cell (i.e., *E. huxleyi* 2090-BD5) within a susceptible population (*E. huxleyi* 2090) in the absence of viral pressure (Fig. 3D) implies that heterogeneity within the susceptible population can prevent its extinction by EhV infection. These resistant cells can survive viral infection to form a new population (Fig. 1B).

We suggest that the continuous formation of heterogeneous cells with different resistance levels also caused stable *E. huxleyi*-EhV coexistence in the recovered population (2090-Rec). Due to constant formation and lysis of a minority of susceptible cells (Fig. 2A), the virus propagates while the majority of resistant cells continue to multiply. Every division is generating a new generation of heterogeneous cells that includes a new fraction of susceptible cells. Interestingly, no susceptible cells were isolated from the coexisting 2090-Rec population (Fig. 3D), suggesting that susceptible cells are lysed before or after isolation thus preventing the formation of a monoculture. Furthermore, none of the monocultures produced viruses, indicating that all the 2090-Rec derived single cells that yielded monocultures were resistant to the virus upon sorting. These results highlight the necessity of an assortment of cells with susceptible and resistant phenotypes for a lytic virus to coexist with its host. Furthermore, the screening for resistance of a population derived from a single cell (Supplementary Fig. S7) implies that individual cells can develop phenotypic heterogeneity within only a few generations. All cells within a monoculture share the same genetic background, pointing out that virus resistance may be a highly plastic trait.

That cell heterogeneity can facilitate the coexistence between a host and a lytic virus has been observed in various other microbial systems. An early observation by Delbruck et al. pointed out that “reverse mutations” can cause a small number of bacterial cells to become susceptible to phage infection (39). The observation that viral infection during host-virus coexistence is limited to a subpopulation of susceptible cells was also found in the marine microalga *Ostreococcus mediterraneus* (6), the marine cyanobacterium *Prochlorococcus* sp. (40), and *Escherichia coli* (9), with the underlying mechanism having been attributed to horizontal gene transfer, gene mutations and genome rearrangements. The source of the observed cell-to-cell heterogeneity in the coexisting *E. huxleyi* population remains elusive. Possible explanations for the emerged resistant phenotypes may involve (i) genetic modifications, including random mutations (41), chromosome rearrangement (42), mobile genetic elements such as transposons (43, 44) and horizontal gene transfer (45), as well as (ii) non-genetic modifications, such as methylations that impact epigenetics (46, 47) or programmed transcriptional responses (28, 48). The isolation of monoculture 2090-BD5 from an uninfected *E. huxleyi* 2090 culture indicates spontaneous changes can lead to a resistant phenotype. In addition to generating cells with heterogeneous resistant levels, previous reports suggested that coexistence on the population level can be mediated by molecules released during cell lysis that prevent virus attachment to *E. huxleyi* (21). Lastly, another mechanism might be the concept of a “numerical refuge” where low densities of susceptible cells avoid extinction through a low probability of virus encounter (49).

### Continuum of virus resistant phenotypes as a bet-hedging strategy

By mapping virus resistance within a host population at single-cell resolution, we revealed a spectrum of resistance levels within *E. huxleyi* cultures (Fig. 3). As described by the ‘Kill the Winner’ hypothesis, virus resistance comes with a cost (50), such as lower nutrient assimilation (9, 41), higher susceptibility to viral infection by other viruses (40, 45, 51), or reduced growth rate (8, 26, 52), as was observed also in this study (Supplementary Fig. S10). Thus, the high cell-to-cell heterogeneity may provide a bet-hedging strategy to cope with changing environments (53). By maintaining a reservoir of diverse cell types with different degrees of susceptibility, the environment will select for the best-fitted ones, eventually leading to variations in the phenotypic composition among populations. The many levels of resistance are a suitable strategy to optimize the tradeoff between virus resistance and varying levels of environmental stresses and host physiological states. We hypothesize that the presence or absence of viruses shapes the population composition by promoting either resistant, less competitive (lower µ), or susceptible, highly competitive cells, respectively. We suggest that the selection pressure of the environment is the guiding principle of the dissimilar population composition of the naïve *E. huxleyi* 2090 population compared to the coexisting *E. huxleyi* 2090-Rec population. For example, the naïve *E. huxleyi* 2090 population consists of a majority of susceptible cells (Fig. 3B), while the *E. huxleyi* 2090-Rec population consists of a majority of resistant cells (Fig. 3D, Fig. 2A). The resistant *E. huxleyi* 2090-BD5 strain, which was isolated from an uninfected *E. huxleyi* 2090 population, grows slower than *E. huxleyi* 2090 when co-cultured in the absence of viruses (Supplementary Fig. S12F), illustrating how susceptible cells can take over the population without viral pressure. Furthermore, in the absence of viruses, resistant monocultures occasionally displayed a loss of the resistant phenotype over several months (data not shown). Our results suggest that susceptible cells have an advantage in virus-free environments (Supplementary Fig. S12F), while cells with a high resistance level have an advantage in an environment with a high viral load (Fig. 4B, C). Importantly, producing a heterogeneous population with a fraction of susceptible cells possesses an additional advantage. The production of susceptible cells supports EhV proliferation, which may lyse co-occurring susceptible strains providing an effective weapon of infectious virions against other *E. huxleyi* susceptible strains that are evolving during bloom succession. This intraspecies competition via viral production has been suggested in bacteria and archaea (54–56), however, such an interaction between eukaryotes and lytic giant viruses was never reported (Fig. 4A). The cost for *E. huxleyi* coexistence with EhV is the death of the susceptible fraction of the population and proliferation of only the resistant cells, and the risk of competing against a strain with a higher resistance level (2090-Rec *vs*. 2090-BD5 and 2090-Rec vs. 379). The *E. huxleyi*-EhV coexistence simultaneously demonstrates the predation of some cells with beneficial consequences for the entire population.

The finding of multiple resistance phenotypes stretching on a continuum (57, 58) offers a novel perspective on the binary classification of susceptibility or resistance. This continuum adds complexity to the species or strain-specific network of host-virus interactions. Our work unveils a higher resolution of host specificity and implies that the infection outcome may vary depending on the individual cell. The cellular mechanisms that can account for different levels of resistance against viral infection within *E. huxleyi* populations are currently unknown and may comprise various strategies. Previous studies indicated that virus resistance can be associated with morphological changes in the formation of organic scales over the *E. huxleyi* cell surface (26) and unique lipid composition (36). The recently identified lipid biomarkers for resistance were not identified in resistant cultures of *E. huxleyi* 2090-BD5 and 2090-Rec (data not shown), suggesting that their resistance did not arise from membrane alterations. Gaining virus resistance may also be tightly linked to cell metabolism. Viral infection is highly affected by the cellular physiological state due to the dependency on cellular metabolic pathways, cell physiology (31) and resources for viral progeny (38). As a giant virus with a large burst size (59–61), EhV has a high metabolic demand that must be met by the host cell. This offers numerous possibilities by which remodeling of host metabolism can lead to varying levels of cell resistance with metabolic costs. Differential expression of metabolic genes was shown to be associated with resistance or susceptibility in various *E. huxleyi* strains (36, 61, 62). Lastly, various defense mechanisms may lead to resistance, as was found in prokaryote genomes that can carry a wide arsenal of anti-viral systems. Utilization of more than one defense system often has a complementary, overlapping, or cumulative effect (63) against different viral species. Further investigation is needed to identify anti-viral defense mechanisms in *E. huxleyi* and their distribution among cells in the population to explain the continuum of resistance levels.

### Ecological significance of E. huxleyi’s phenotypic heterogeneity

The genetic diversity of *E. huxleyi* and EhV strains reported in natural blooms (64, 65) highlights the complexity of *E. huxleyi* -EhV interactions in the ocean. We used a simple model of an *E. huxleyi* culture and its derived clones as an experimental model system to reduce the complexity of natural *E. huxleyi* community and ask fundamental ecological questions regarding host-virus coexistence.

Based on the isolation of an innate resistant cell (Fig. 3D), it is likely that bloom termination by EhV induces a strong selection pressure for the survival of rare pre-existing resistant *E. huxleyi* subpopulations during bloom demise and act as a seed population within the bloom for subsequent blooms. Although further quantification of the fraction of infected and resistant cells is required during and after bloom events, our results suggest that following bloom demise, resistant cells may form coexistence with the lytic EhV by generating a new susceptible daughter population. We hypothesize that recovery of the resistant population after bloom termination by EhV may occur similarly to the recovery observed in the lab (Fig. 1B, Supplementary Fig. S13). However, bottom-up conditions and other mortality agents of *E. huxleyi* probably limit the growth of the new population. Under the condition of sufficient host-virus contact rates, the coexistence of *E. huxleyi* and EhV could benefit both the alga and the virus by increasing algal competitiveness and prolonged association of EhV with its host, particularly during the long periods in-between annual blooms. However, if the probability of virus encounters is extremely low, our findings indicate that in the absence of viral pressure the cells which are typically bloom-forming; opportunistic, competitive, and virus-susceptible (27, 66) will outcompete the slow growing resistant cells (Supplementary Fig. S12F). This illustrates that a population of resistant cells that survived a virus attack could be outgrown by susceptible cells setting the stage for the next bloom event. This observation may contribute to previous reports showing high seasonal stabilities of marine microbial and viral communities (67, 68) and the reoccurrence of the same *E. huxleyi* and EhV genotypes in a multi-annual manner (69).

The use of single-cell analysis to characterize virus resistance at single-cell resolution revealed a large cell-to-cell heterogeneity. Studying *E. huxleyi* phenotypes by comparing cells with identical genetic backgrounds and distinct resistant phenotypes possesses a potential for new discoveries of ecologically relevant anti-viral strategies. We propose that the generation of phenotypic heterogeneity is pivotal for the *E. huxleyi*-EhV arms race in the natural environment and allows the emergence of better adapted cells to survive. Heterogeneity is thus a driving force to enable the coexistence of *E. huxleyi* and EhV.

## Material and methods

### Strains of E. huxleyi and EhV

The following *E. huxleyi* strains were obtained from the Roscoff Culture Collection (RCC) or from the National Center for Marine Algae and Microbiota (NCMA): CCMP2090, CCMP373, CCMP374, CCMP379, and RCC1216 (hereinafter: *E. huxleyi* 2090, 373, 374, 379, and 1216 respectively), as well as RCC9645, RCC9646, and RCC9655, which were isolated in 2018 from a mesocosm experiment in Bergen (Norway). The *E. huxleyi* strains Rec-17, Rec-32, Rec-53, and Rec-97 were derived by single-cell isolation from the recovered coexisting culture *E. huxleyi* 2090-Rec. The *E. huxleyi* strains 2090-BD5 and 2090-2 were derived by single-cell isolation from *E. huxleyi* 2090. For viral inoculation, the following EhV strains were used: EhV-201, EhV-86, EhV-163 (14), EhV-ice (15), EhV-M1 (70), and EhV-Rec and propagated on *E. huxleyi* as listed in Supplementary Table S2.

### Culture maintenance and experimental conditions

Algal cultures were kept at 18°C with a 16:8 hours light:dark cycle and a light intensity of 100 μmol photons m^-2^ s^-1^, provided by cool white light-emitting diodes. Cultures were diluted weekly at a ratio of 1:10 into fresh medium. The medium was composed of autoclaved and filtered seawater (FSW) supplemented with modified K/2 medium (replacement of organic phosphate with 18 μM KH_2_PO_4_)(Gerecht et al., 2014)(Gerecht et al., 2014)) and the antibiotics ampicillin (100 μg mL^-1^) and kanamycin (50 μg mL^-1^). For regular virus infection assays, algal cultures were grown in 50 mL tissue culture flasks. For infection assays that included smFISH sampling, cultures were grown in 650 mL tissue culture flasks.

### Viral lysate preparation and algal culture inoculation

A fresh viral lysate was propagated one week prior to every infection assay. Each EhV strain was propagated with its respective algal host strain (Supplementary Table S2). In brief, 1 L of an exponentially growing *E. huxleyi* culture at 1-2×10^6^ cells mL^-1^ was inoculated with EhV in a 5:1 virus-to-cell ratio. After 4 days, the viral lysate was filtered through a 1.2 µm pore size glass microfiber filter (grade GF/C, GE Healthcare Whatman) followed by a 0.45 µm pore size Nalgene Rapid-Flow filter unit (PES, ThermoFisher Scientific) to remove cell debris, before concentrating and washing the virions by 100 kDa tangential flow filtration (Vivaflow 200, Sartorius). The concentrated viral lysates were sterile-filtered through a 0.22 µm filter (PVDF, Millex-GV, Millipore) and stored in darkness at 4°C until the infection assay. For EhV-Rec, which co-exists with the algal host *E. huxleyi* 2090-Rec, no inoculation was needed. EhV-Rec virions were filtered and concentrated as described above, however, final filtration through 0.22 µm was avoided due to the loss of virions possibly due to the formation of transparent exopolymer particles. For every infection assay, an exponentially growing algal culture at ∼5x10^5^ cells mL^-1^ was inoculated at a 1:1 virus-to-cell ratio.

### Enumeration of algal cell abundance, cell death and viral abundance

Algal cells and viruses were monitored by flow cytometry (Supplementary text). In brief, living algal cells identified by their chlorophyll autofluorescence, while the fraction of dead algal cells was quantified following the staining with Sytox Green (Supplementary Fig. S14A-C). Viral particles were fixed with glutaraldehyde and stained with SYBR Gold before flow cytometry analysis (Supplementary Fig. S14D). Data analysis was conducted using CytExpert 2.4 (Beckman Coulter).

### Calculation of growth rate (μ), carrying capacity (CC) and maximum viral production (MVP)

The growth rate (μ) was calculated as *μ* = ln(*N*_2_/*N*_1_)/ (*t*_2_ − *t*_1_), where *N*_*1*_ represents the cell abundance at time 1 (*t*_*1*_), and *N*_*2*_ represents the cell abundance at time 2 (*t*_*2*_) (72). *t*_*1*_ is the first day of the exponential growth phase of the untreated cultures, and t_2_ is the last day that a culture grows exponentially. The same calculation was conducted to estimate *μ* for the EhV-inoculated cultures. The carrying capacity (CC) is the highest cell abundance that each *E. huxleyi* strain reached in stationary phase. The maximum viral production (MVP) represents the highest abundance of extracellular viral particles as reached in stationary phase cultures. All measurements were conducted for EhV-inoculated and non-inoculated cultures of each *E. huxleyi* strain in three biological replicates.

### Quantification of actively infected E. huxleyi cells using smFISH

To estimate the fraction of infected cells within an *E. huxleyi* population, we used smFISH based on the EhV *mcp* gene, as previously described (4). In brief, a mix of 47 probes of 20 nucleotides was designed to bind the *mcp* mRNA of EhV at different locations. Conjugation with the fluorophore tetramethylrhodamine (TMR) allows the detection by flow cytometry (ex: 561 nm, em: 564-606 nm). Cells with >10^3^ A.U. area were enumerated as *mcp* positive cells. Aliquots of 30 mL algal culture were fixed with paraformaldehyde (1% final concentration) and incubated for 1 h at 4°C with agitation. Samples were centrifuged for 2 min, the supernatant was discarded, and the pellets were re-suspended in 1 mL cryopreserving solution. To ensure contact of the cryopreservant with all fixed cells before freezing, samples were incubated for 1 h at 4°C with agitation. Samples were centrifuged, the supernatant removed, and the pellet stored at -80°C until hybridization. For flow cytometry analysis, samples were thawed at RT, and chlorophyll was extracted by sequential re-suspension in 70% and 90% ethanol (HPLC grade, J.T. Baker). Samples were treated with 500 μL proteinase-K (10 μg mL^-1^ final concentration, Ambion). For hybridization, 50 μL of hybridization buffer (17.5% formamide concentration) containing the *mcp* probes (0.1 ng mL^-1^ final concentration) were added to all samples for overnight incubation at 30°C in darkness. Lastly, samples were stained with the nucleic acid stain DAPI (10 μg mL^-1^ final concentration), re-suspended in 400 μL GLOX buffer and analyzed by flow cytometry (FSC height threshold = 5,000 A.U., SSC area threshold = 1,000 A.U.).

### Generation of monocultures by single-cell sorting and screening for viral resistance

To generate clonal cultures (herein monocultures) from heterogeneous *E. huxley*i cultures, single cells were sorted using fluorescence-activated cell sorting (FACS) with BD FACSAria III Cell Sorter (BD Biosciences). Cells were gated based on their optical properties (chlorophyll autofluorescence ∼5×10^4^ A.U., forward scatter ∼5×10^4^ A.U.) and sorted into 200 μL fresh medium in 96-well plates. The well plates were wrapped with parafilm to prevent evaporation and incubated at 18°C and low light (∼20 μmol photons m^-2^ s^-1^) with a 16:8 hours light:dark cycle. After 4-8 weeks, wells that showed algal growth were transferred to new well plates and maintained by a 10-fold dilution into fresh medium every three weeks. Roughly 20% of the sorted cells formed a monoculture that maintained cell growth sufficient for conducting experiments (Supplementary Table S2). To screen for viral resistance, the monocultures were transferred to two 96-well plates, one serving as an uninfected control and the other for inoculation with EhV, and cultured at ∼100 μmol photons m^-2^ s^-1^. Cell growth was monitored daily by measuring chlorophyll autofluorescence in a Tecan Infinite 200 PRO plate reader (ex: 480 nm, em: 660 nm). To convert chlorophyll emission to cell abundances, a late exponential culture was diluted in an 8-step dilution and measured in parallel by flow cytometry and the plate reader, thus, constructing a calibration curve (Supplementary Fig. S15). Monocultures were inoculated at a 1:1 virus-to-cell ratio, according to the average chlorophyll emission of a 96-well plate. At 6 dpi, aliquots of 98 µL from each well were fixed for enumeration of viral particles as described above, thus, ending the experiment.

### Quantification of lipid markers for viral infection using UPLC-HRMS

To assess the occurrence of virus-infected cells within *E. huxleyi* populations by the formation of vGSLs, we used an LC-MS based untargeted lipid profiling approach as previously described (36). In brief, cultures of *E. huxleyi* 2090-Rec, 2090-BD5, and *E. huxleyi* 2090 with and without addition of EhV-201 were analyzed for cellular lipid composition in three technical replicates (except 2090-BD5, for which two replicates were analyzed). Cultures were grown in 1 L FSW + K/2 with ampicillin and kanamycin as described above. Samples of *E. huxleyi* 2090-Rec, 2090-BD5 and 2090 were collected during exponential growth phase, while *E. huxleyi* 2090 inoculated with EhV-201 was collected at 3 dpi, during culture lysis. The samples (150-200 mL of each culture, equivalent to ∼1-3×10^8^ cells per sample) were collected by vacuum filtration onto glass microfiber filters (grade GF/A, GE Healthcare Whatman), immediately plunged into liquid nitrogen, lyophilized to dryness, and stored at -80°C. Lipids were extracted using methyl tert-butyl ether (MTBE) and glucosyl(β) ceramide d18:1/c12:0 as internal standard (Supplementary text). Extracts were dried under a flow of nitrogen (TurboVap, LV, Biotage, Uppsala, Sweden) and stored at -80°C. For UPLC-HRMS analysis, samples were re-dissolved in 200 μL mobile phase B, transferred to glass inserts and an aliquot of 1 µL injected for UPLC-HRMS analysis (Supplementary text). The lipid markers hGSLs, sGSLs and vGSLs were annotated based on their accurate mass and fragmentation pattern (35, 73, 74). Peak areas of indicative masses ([M+Na]^+^ for hGSL and vGSL species, [M+H-Sialic acid]^+^ for sGSL species) above a signal-to-noise threshold of 3 (limit of detection) were normalized to the internal standard and the total number of extracted cells.

### Enumeration of extracellular viruses in E. huxleyi 2090-Rec derived monocultures by qPCR

To verify if the single-cell isolates from *E. huxleyi* 2090-Rec produced viruses, we assessed EhV presence in the extracellular medium of the monocultures derived from *E. huxleyi* 2090-Rec using qPCR and primers for the *mcp* gene of EhV. An aliquot of each monoculture was centrifuged and 50 µL of cell-free supernatant were boiled for 20 min at 100°C to release DNA from virions. Samples were diluted in 120 μL of ultra-pure water to prevent qPCR reaction inhibition by medium substances. The qPCR analysis was conducted as previously described (75), using 5′-acgcaccctcaatgtatggaagg-3′ (mcp1Fw) and 5′-rtscrgccaactcagcagtcgt-3′ (mcp94Rv) as primers. A concentrated EhV-Rec lysate was used as positive control and FSW as negative control. All reactions were carried out in technical triplicates.

### Co-cultivation experiments of two algal strains

To assess the outcome of intraspecies competition, we co-cultured pairs of *E. huxleyi* strains separated by a 0.4 µm membrane, allowing the free exchange of viruses, inorganic nutrients, and metabolites such as vitamins. One algal strain was inoculated into the wells of a 12-well plate and the second strain into Thincert inserts (Greiner). Cell abundances and virus production were monitored for both strains located within each vessel. Before sampling, cultures were re-suspended with a serological pipette to counteract the sedimentation of cells and enhance medium exchange. The pore size of 1 µm allowed viruses to pass the membrane, thus, the enumerated virus abundances cannot be attributed to a single algal strain when both strains were susceptible. Experiments were conducted in three biological replicates. As a control, all strains were co-cultured with themselves by inoculating them both in the wells and in the inserts (Supplementary S13A-E).

### Statistical analysis

The significance of cell abundance and virus abundance of 2090-Rec treated with different EhV strains (Fig. 1D, Supplementary Fig. S3A) was calculated using one-way ANOVA with Šidák correction for multiple comparisons by comparing the mean of each treatment to the non-treated culture at 7 dpi. The statistical significance of the differences in chlorophyll fluorescence and virus production between *E. huxleyi* 2090-Rec-derived monocultures and 2090-derived monocultures (Fig. 3A, B) was assessed using a two-tailed nonparametric Mann-Whitney test\. The coefficient of variance (CV) was calculated as follows: CV = standard deviation/sample mean x 100. The correlation between chlorophyll fluorescence and virus production of *E. huxleyi* 2090-Rec monocultures was calculated by nonparametric Spearman correlation. The significance of the differences in growth rates and carrying capacity across all strains and treatments was estimated using two-way ANOVA with Tukey’s multiple pairwise comparisons. The significance of the differences in maximum viral production was estimated using one-way ANOVA with Tukey’s multiple pairwise. The effect of EhV-Rec on the growth of *E. huxleyi* 2090-BD5 was assessed using a two-tailed t-test to compare the cell abundances of *E. huxleyi* 2090-BD5 co-cultured with *E. huxleyi* 2090-Rec (Fig. 4E) and *E. huxleyi* 2090-BD5 co-cultured with itself (Supplementary Fig. S12C) on day 7 of the co-cultivation. All calculations were conducted using GraphPad Prism 9.3.1 (GraphPad Inc.).

## Acknowledgments

We are grateful to Ron Rotkopf for his help in data analysis, Dr. Flora Vincent for her help in using the smFISH protocol, and Avia Mizrachi for her help during the single-cell sorting. Dr. Daniella Schatz and Ben Labbel helped us with experimental design and data analysis. This research was supported by the Simons Foundation grant (no. 735079) “Untangling the infection out-come of host-virus dynamics in algal blooms in the ocean” awarded to A.V.

## Author contributions

N.J., C.K., G.S. and A.V. conceptualized the study and experimental design; N.J. and N.S. performed the research; C.K. performed the lipid extraction and analysis; N.J., C.K. and A.V. wrote the manuscript; G.S. and A.S. reviewed and edited the manuscript.

