## Supplementary data for "Cell-to-cell heterogeneity drives host-virus coexistence in a bloom-forming alga"

**This PDF file includes:**

Supporting text
Figures S1 to S15
Tables S1 to S2

**Supplementary material and methods**

*Enumeration of algal cell abundance, cell death and viral abundance using flow cytometry*

All flow cytometry analyses were performed using a CytoFLEX S Flow Cytometer (Beckman Coulter, Nyon, Switzerland). Living algal cells were identified by plotting the chlorophyll autofluorescence (excitation (ex) at 561 nm, emission (em) at 665-715 nm) versus the forward scatter (FSC) as a proxy for cell size (FSC area threshold = 90,000 arbitrary units (A.U.)). Only cells with an autofluorescence >$4.5\times10^4$ A.U. were enumerated as live cells (Supplementary Fig. S14A). To quantify dead algal cells, we evaluated the percentage of cells that were stained by Sytox Green (Invitrogen, Supplementary Fig. S14B) based on the increased membrane permeability of dead cells. Samples were stained at a final concentration of 1 µM Sytox Green and incubated in the dark for 30 min at room temperature (RT) before flow cytometry analysis (ex: 488 nm, em: 505-545 nm). Unstained samples were used as control to correct for background fluorescence (Supplementary Fig. S14C). Per sample, 200-200,000 events of low and high chlorophyll emitting cells were enumerated. To quantify extracellular viral particles, samples were fixed with glutaraldehyde (Sigma) at 0.5% final concentration, incubated for 30 min at 4°C, and frozen with liquid nitrogen before storage at -20°C. Fixed samples were thawed and stained with the nucleic acid stain SYBR Gold (Invitrogen), and then diluted 1:10,000 in Tris-EDTA buffer. The samples were incubated for 20 min at 80°C and cooled to RT before flow cytometry analysis (ex: 488 nm, em: 500-550 nm; Violet SSC area and FITC area threshold = 1,000 A.U.). Per sample, 3,000-300,000 events were enumerated together with the estimation of the background noise (Supplementary Fig. S14D).

*Quantification of lipid markers for viral infection using UPLC-HRMS*

Lipid extraction and UPLC-HRMS analysis were performed as previously described (36). In brief, 3 mL of a pre-cooled (-20°C) methanol:methyl tert-butyl ether (MTBE) (1:3, v:v) solution was added to each filter that contained a sphingolipid standard mixture with 146 nM of the glucosyl(β) ceramide d18:1/c12:0, which was used as internal standard. The samples were shaken, sonicated and supplemented with 1.5 mL water:methanol (3:1, v:v) to obtain phase separation upon centrifugation. The upper organic phase (1.5 mL) was transferred to 2 mL centrifuge tubes and dried under a flow of nitrogen. The lower polar phase was re-extracted with 1.5 mL of MTBE. The upper organic phase (2.5 mL) was combined with the organic phase from the first extraction and dried under a flow of nitrogen (TurboVap, LV, Biotage, Uppsala, Sweden). The samples were stored at -80°C until UPLC-HRMS analysis. One blank filter was processed as extraction blank.

For UPLC-HRMS analysis, samples were thawed, re-dissolved in 200 µL acetonitrile:isopropanol (7:3) with 1% 1 M ammonium acetate and 0.1% acetic acid, transferred to 200 µL glass inserts in autosampler vials and directly used for injection. An aliquot of 1 µL was analyzed using UPLC coupled to a photodiode detector (ACQUITY UPLC I-Class, Waters, Milford, MA) and a quadrupole time-of-flight mass spectrometer (SYNAPT G2 HDMS, Waters). Full scan and all ion fragmentation ($MS^E$) analysis were performed in positive ionization mode alternating with 0.1 min scan time between low (1 eV) and high (ramp of 15-35 eV) collision energy.

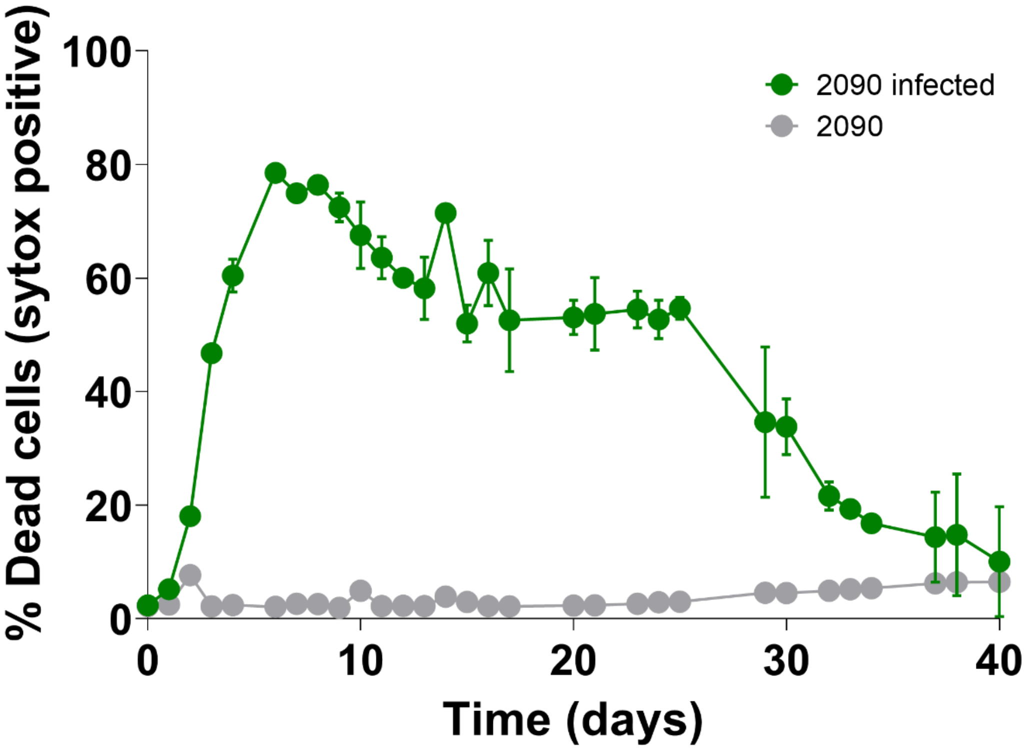


**Figure S1. Cell death in *E. huxleyi* 2090 cultures with and without EhV-201 inoculation.** Dead cells were enumerated using the cell death marker Sytox Green. Values are presented as mean ± SD (n = 3).

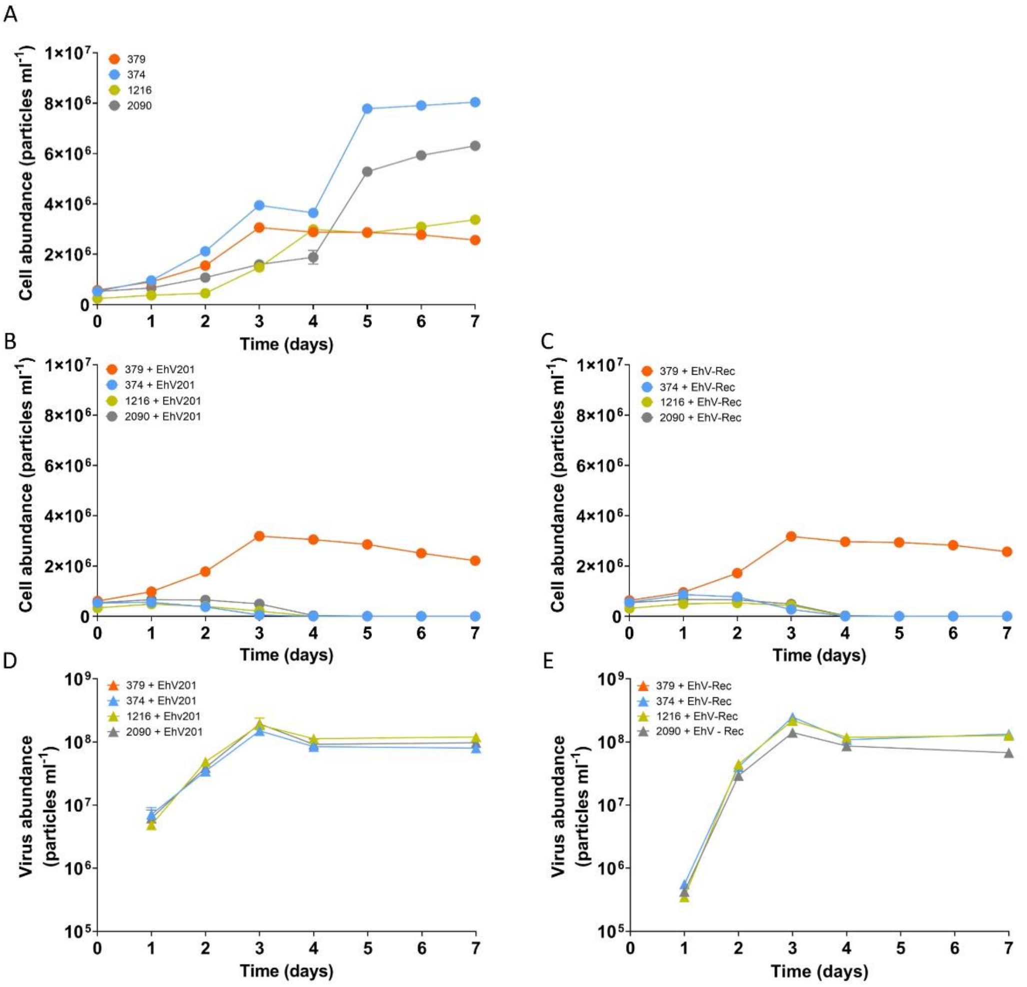


**Figure S2. Infection dynamics of EhV-Rec and its ancestor EhV-201 with the *E. huxleyi* strains 379, 374, 1216 and 2090.** (A) Cell abundances of non-treated cultures. (B) Cell abundances of the cultures inoculated with EhV-201. (C) Cell abundances of the cultures inoculated with EhV-Rec. (D) Extracellular viruses in EhV-201-treated cultures. (E) Extracellular viruses in EhV-Rec-treated cultures. No virus particles were detected in the non-inoculated control cultures and in virus-inoculated cultures of strain 379. Values are presented as mean ± SD (n = 3).

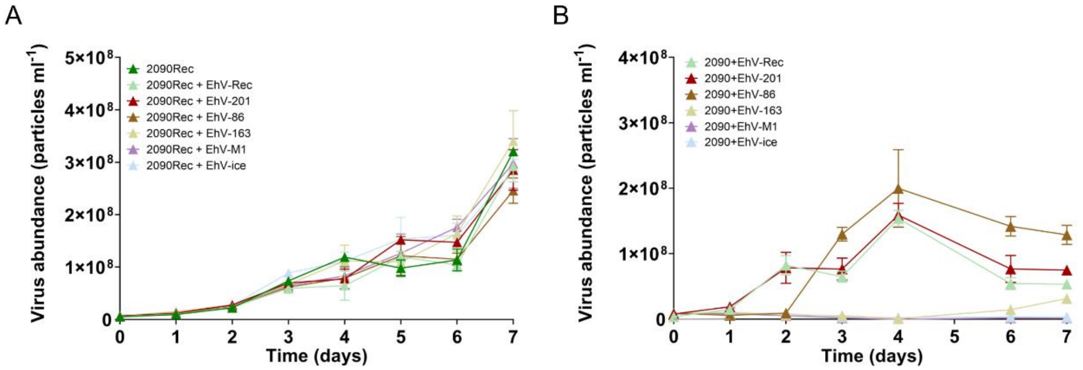


**Figure S3. Virus production in *E. huxleyi* 2090-Rec and its ancestor *E. huxleyi* 2090 following inoculation with several EhV strains that vary in their infectivity.** Each of the two algal strains was inoculated separately with EhV-Rec, EhV-201, EhV-86, EhV-163, EhV-M1 and EhV-ice. (A) Virus production in *E. huxleyi* 2090-Rec cultures following virus addition. The viruses inoculated with *E. huxleyi* 2090-Rec were added in addition to the constant production of EhV-Rec by the culture. (B) Virus production in *E. huxleyi* 2090 cultures following virus addition. Values are presented as mean ± SD (n = 3).

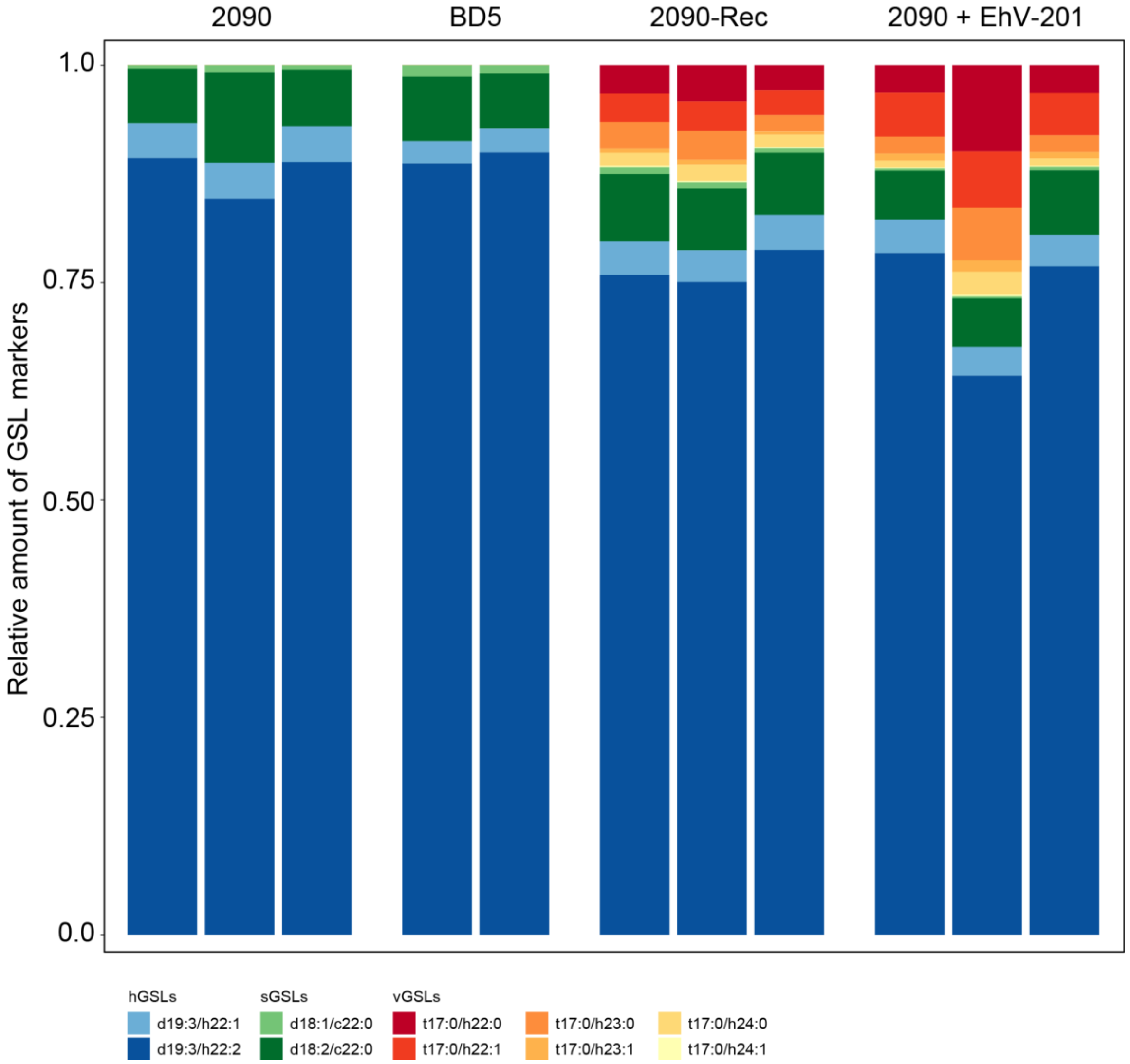


**Figure S4. Relative cell amount of glycosphingolipid (GSL) markers in *E. huxleyi* strains that vary in their host-virus interaction.** Relative cellular amounts of hGSLs (present in all *E. huxleyi* cells), sGSLs (present in susceptible *E. huxleyi* cells), and vGSLs (present in virus-infected cells) in exponentially growing cultures are presented for three *E. huxleyi* strains: the susceptible *E. huxleyi* 2090, the resistant 2090-BD5 (isolated as innate resistant strain from *E. huxleyi* 2090), and the resistant 2090-Rec (host-virus co-existing culture that was recovered following infection of *E. huxleyi* 2090 with EhV-201). Virus-inoculated cultures of *E. huxleyi* 2090 were sampled at 3 dpi. Cultures were sampled in three (2090, 2090+EhV-201, 2090-Rec) or two (2090-BD5) technical replicates.

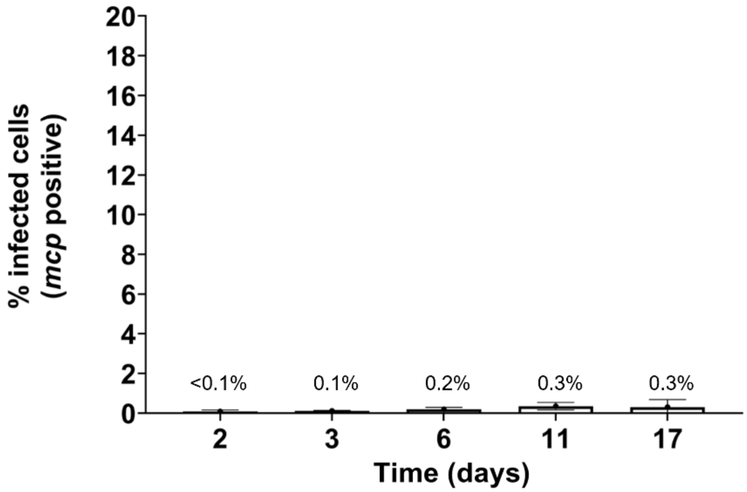


**Figure S5. Fraction of infected cells in uninfected *E. huxleyi* 2090 as quantified by smFISH.** The uninfected *E. huxleyi* 2090 cultures presented in Fig. 1B were sampled five times throughout culture growth. No virus-infected cells expressing the viral *mcp* gene were detected for these cultures (observed percentages <0.5% are within technical noise level). Values are presented as mean ± SD (n = 3).

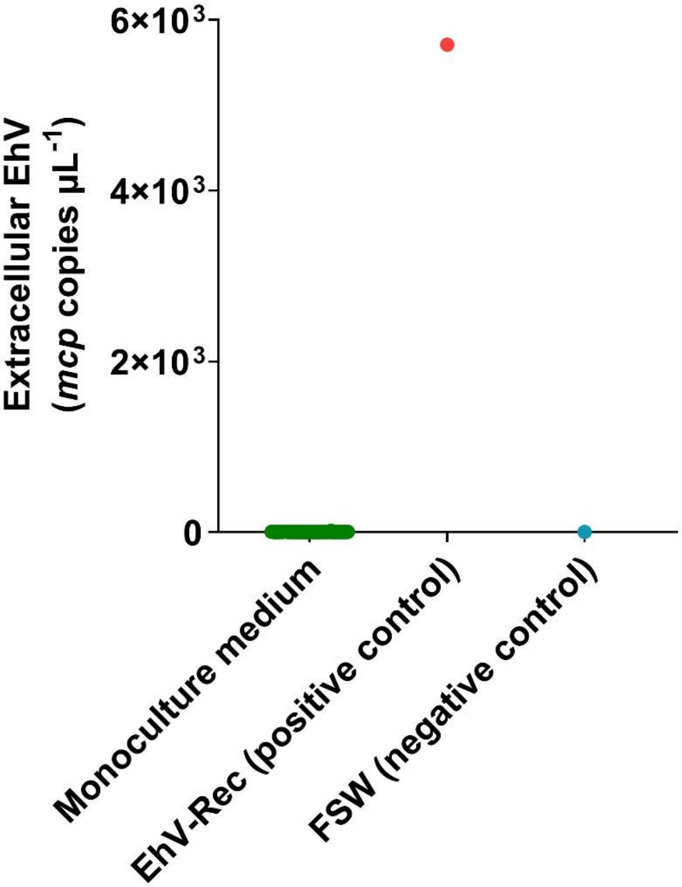


**Figure S6. qPCR amplification of the major capsid protein (*mcp*) gene in the extracellular medium of *E. huxleyi* 2090-Rec-derived monocultures.** To check for production of EhV by monocultures derived from *E. huxleyi* 2090-Rec, we subjected the medium from each monoculture to a qPCR analysis targeting the EhV *mcp* gene marker. The dot plot displays the levels of extracellular virions found in the monoculture medium (●). As a positive control, we processed a concentrated viral lysate from the *E. huxleyi* 2090-Rec culture (●), while filtered seawater (FSW) served as virus-free negative control (●).

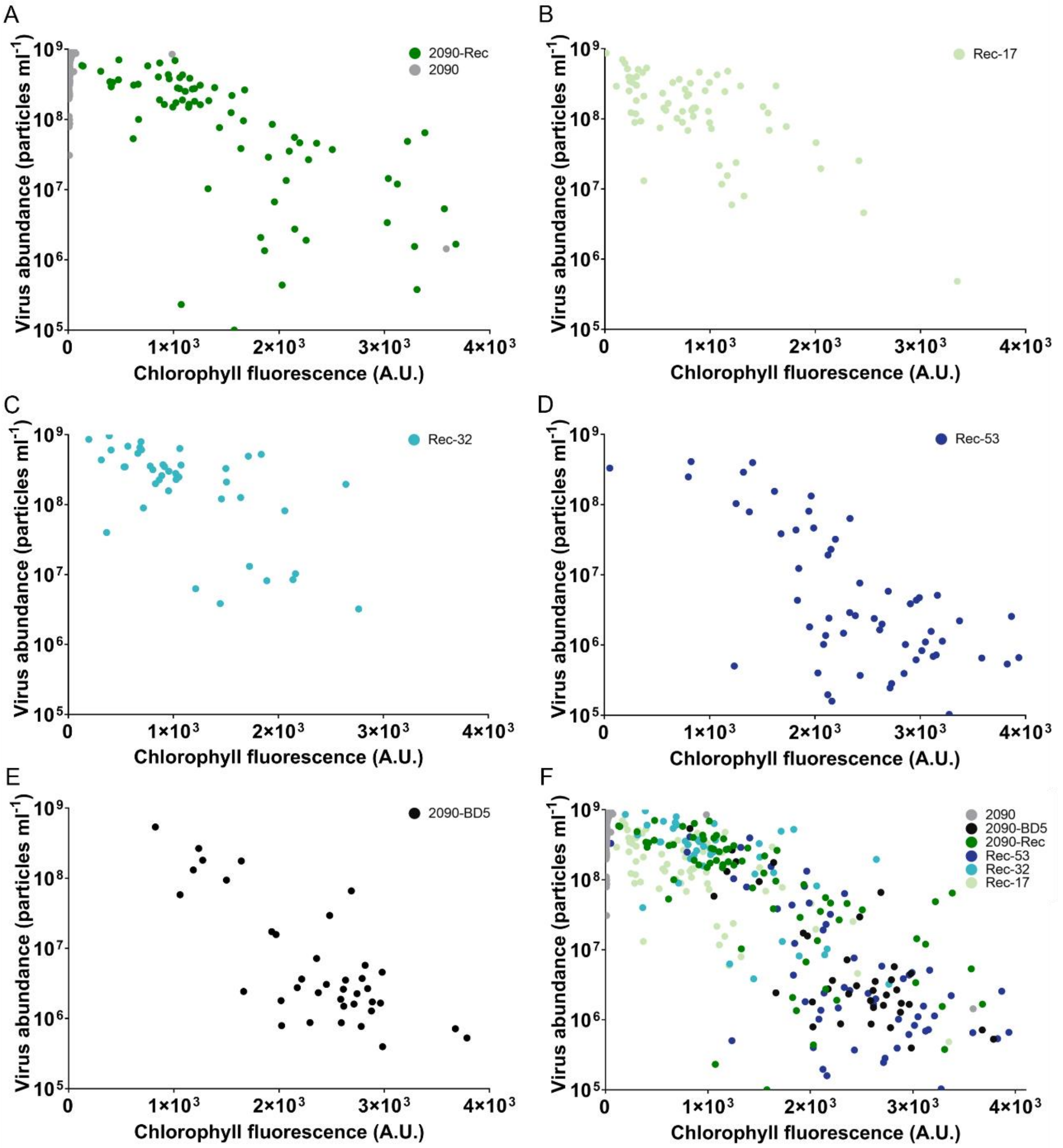


**Figure S7. Monoculture screening for resistance reveals phenotypic heterogeneity with correspondence to the resistant phenotype of the original population.** Monocultures were generated by single-cell sorting and their chlorophyll fluorescence and viral abundance measured at 6 dpi following inoculation with EhV-201. Monocultures originated from the *E. huxleyi* cultures (A) 2090 (*n*=123) and 2090-Rec (*n*=74). From these, monocultures with representative phenotypes were selected for subsequent single-cell sorting and resistance screening, namely (B) Rec-17 (*n*=76), (C) Rec-32 (*n*=43), (D) Rec-53 (*n*=58), and (E) 2090-BD5 (*n*=38). (F) Distribution of all monocultures along the continuum from resistance to susceptibility.

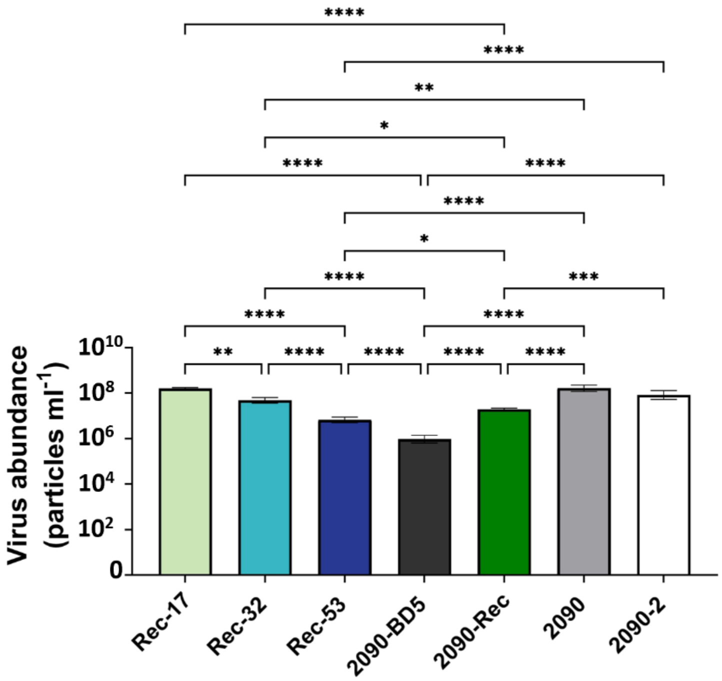


**Figure S8. Maximum virus production as a parameter to define the level of virus resistance.** The maximum value of virus abundance measured in the virus-inoculated cultures throughout 10 days of growth. Average and standard deviation of three biological replicates are shown; * *p*-value < 0.05; ** *p*-value < 0.01; *** *p*-value < 0.001; **** *p*-value < 0.0001. Non-significance is not shown.

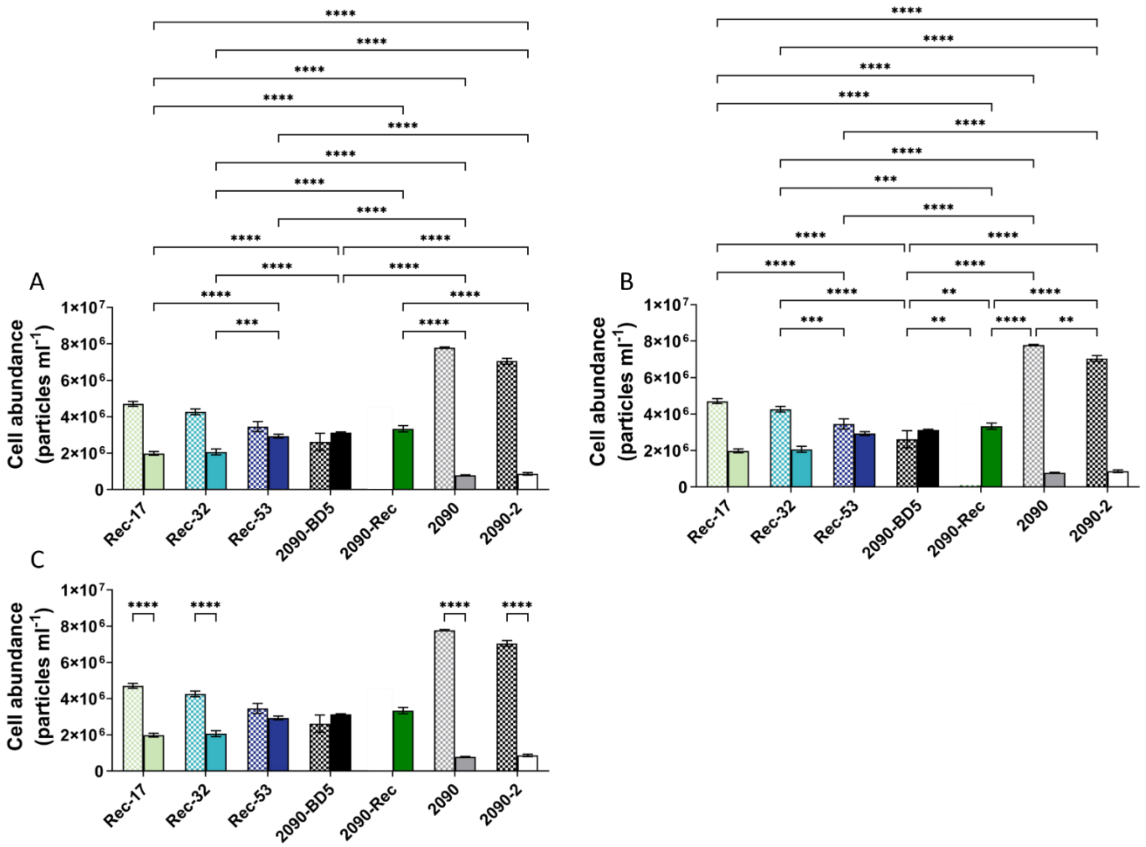


**Figure S9. Maximum carrying capacity as a parameter to define the level of virus resistance.** The maximum cell abundance measured. (A) Pairwise comparisons between virus-inoculated cultures (full filling pattern). (B) Pairwise comparisons between non-inoculated cultures (gridded filling pattern). (C) Pairwise comparisons of virus-inoculated and non-inoculated cultures of the same strain. Average and standard deviation of three biological replicates are shown; * $p$-value < 0.05; ** $p$-value < 0.01; *** $p$-value < 0.001; **** $p$-value < 0.0001. Non-significance is not shown.

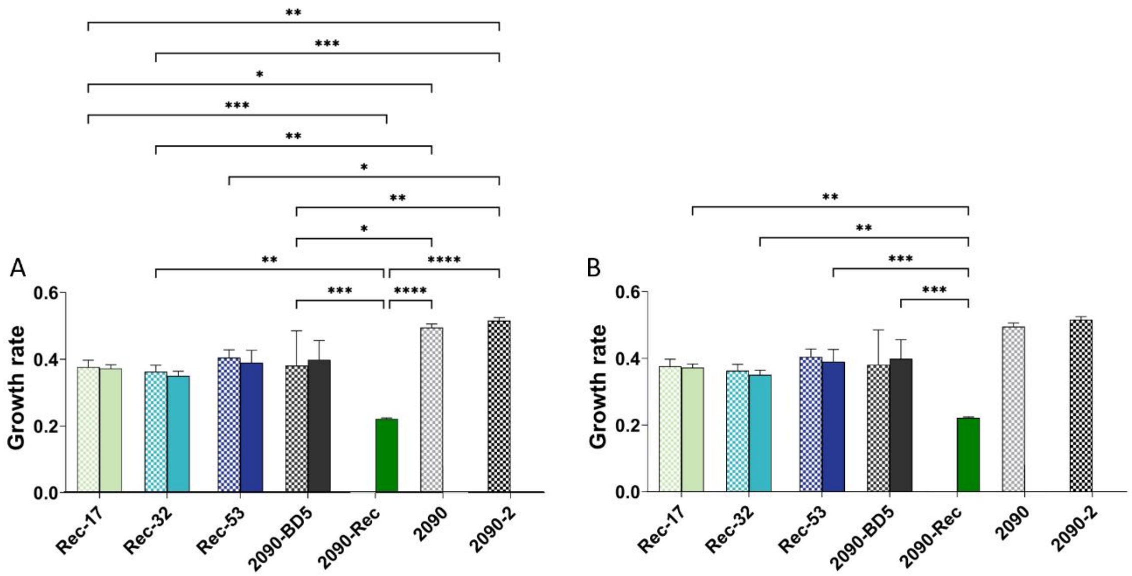


**Figure S10. Growth rate as a parameter to define the level of virus resistance.** (A) Pairwise comparisons between non-inoculated *E. huxleyi* cultures (gridded filling pattern). (B) Pairwise comparisons between virus-inoculated cultures (full filling pattern). Average and standard deviation of three biological replicates are shown; * *p*-value < 0.05; ** *p*-value < 0.01; *** *p*-value < 0.001; **** *p*-value < 0.0001. Non-significance is not shown.

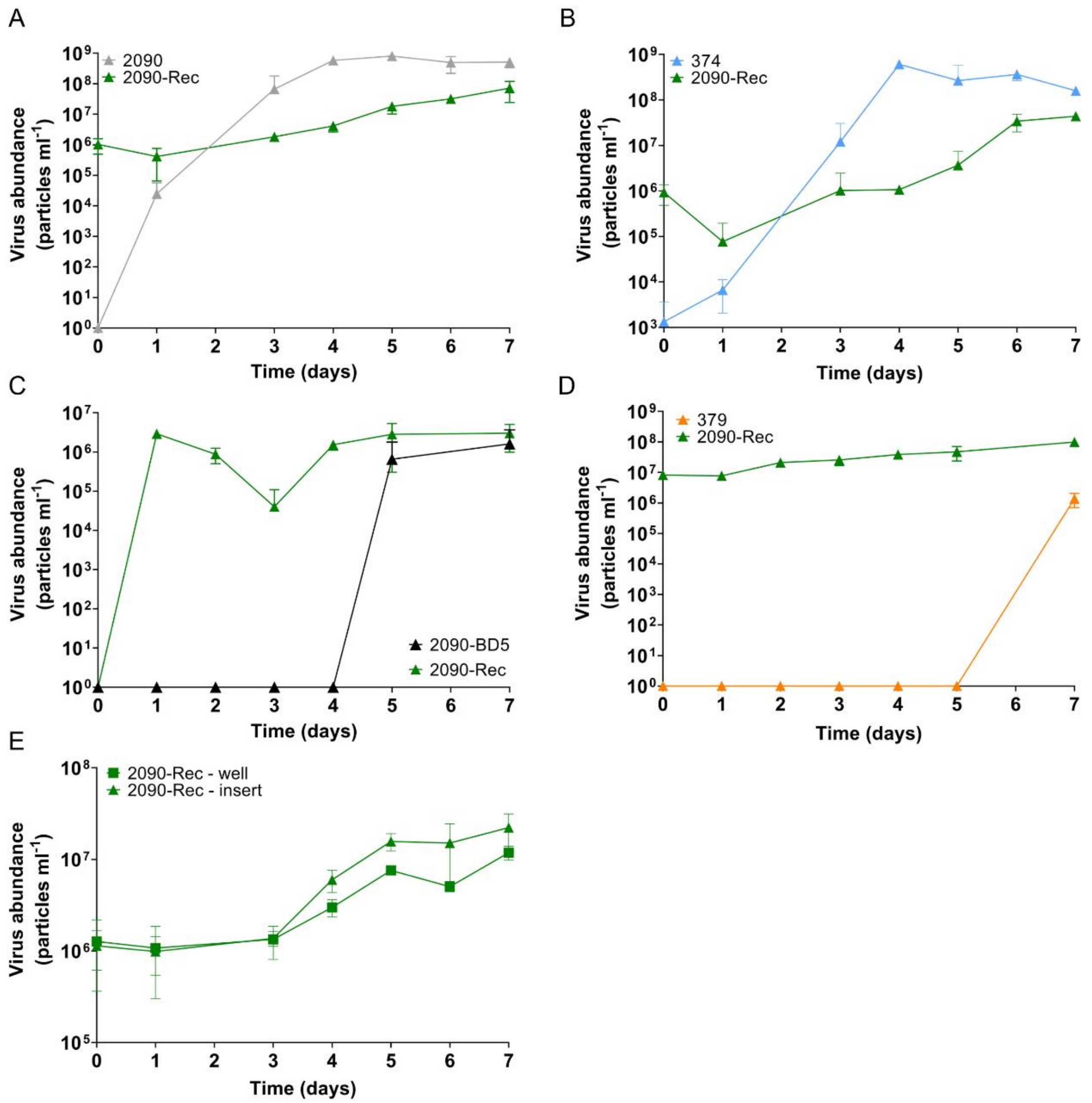


**Figure S11. Virus abundances during co-culturing of 2090-Rec with selected *E. huxleyi* strains that vary in their resistance.** Virus abundances measured during cultivation of the coexisting *E. huxleyi* 2090-Rec with (A) 2090, (B) 374, (C) 2090-BD5, (D) 379, and as a control (E) itself, 2090-Rec. Noteworthy, viruses can pass through the insert membrane which hinders the identification of the strain responsible for virus production. *E. huxleyi* strains were co-cultured in 24-well plates separated by a 1 µm membrane, allowing the exchange of nutrients, metabolites, and viruses, while preventing transfer of algal cells. *E. huxleyi* 2090-Rec was grown in the insert and the competing strains were grown within the surrounding wells. Values are presented as mean ± SD (n = 3).

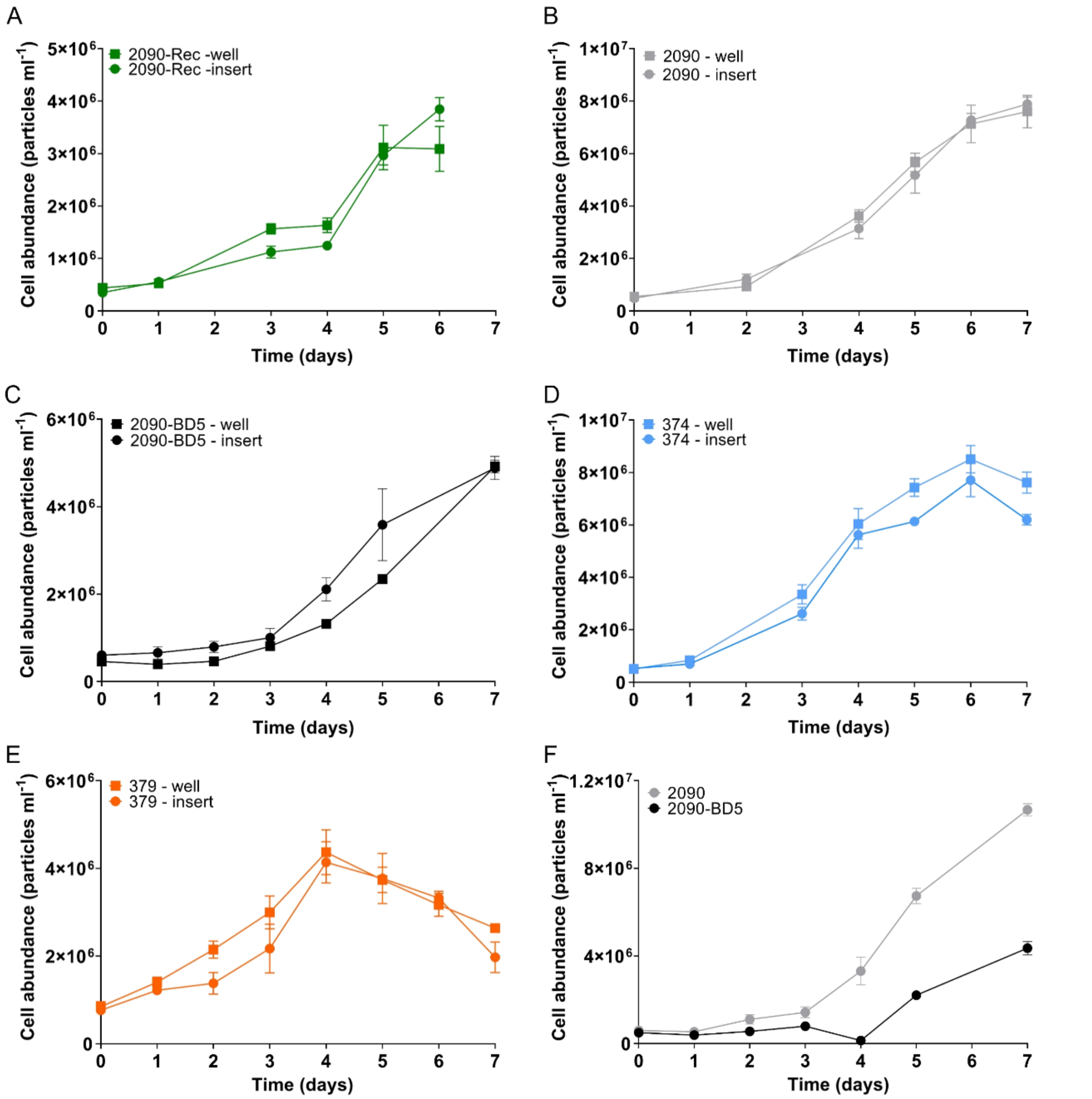


**Figure S12. Cell abundances of selected *E. huxleyi* strains that vary in their resistance when co-cultured with themselves.** *E. huxleyi* strains were co-cultured in 24-well plates separated by a 1 µm membrane, allowing the exchange of nutrients, metabolites, and viruses, while preventing transfer of algal cells. To assess differences in irradiance, strains were inoculated both into the well and into the insert. Cell abundances of (A) *E. huxleyi* 2090-Rec, (B) 2090, (C) 2090-BD5, (D) 374, and (E) 379 show no growth differences between the inserts and wells. (F) Co-cultivation of the susceptible ancestor *E. huxleyi* 2090 with the resistant isolate 2090-BD5. Values are presented as mean ± SD (n = 3).

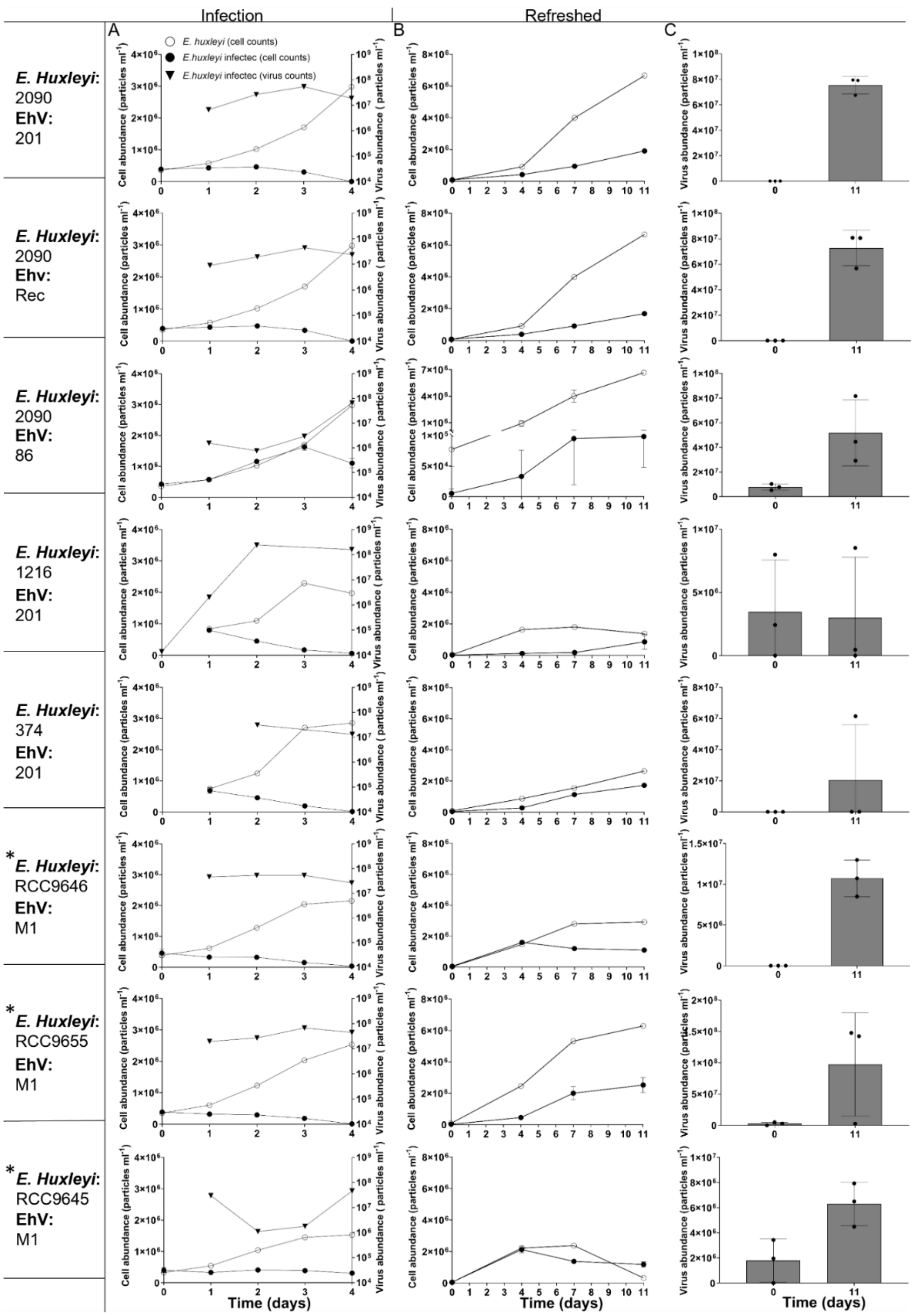


**Figure S13. Host-virus coexistence emerges in a variety of host and virus strains including well-established and recently-isolated *E. huxleyi* strains.** (A) Infection assays with different strains of *E. huxleyi* and lytic EhV led to the population demise and increase in viral particles. (B) Proliferation of recovered, resistant host cells in virus-inoculated cultures that were diluted in fresh growth medium (1:10 *v*:*v*) on 48 dpi of the experiment. For comparison, cell abundances of refreshed non-inoculated cultures are shown. (C) Concomitant proliferation of virions at days 0 and 11 after diluting the virus-inoculated cultures in fresh growth medium. Cell abundances of inoculated cultures (●), cell abundances of non-inoculated cultures (○), virus abundances of inoculated cultures (▼). *E. huxleyi* strains that were isolated in 2018 from induced blooms in a mesocosm experiment (*). Values are presented as mean ± SD (n = 3).

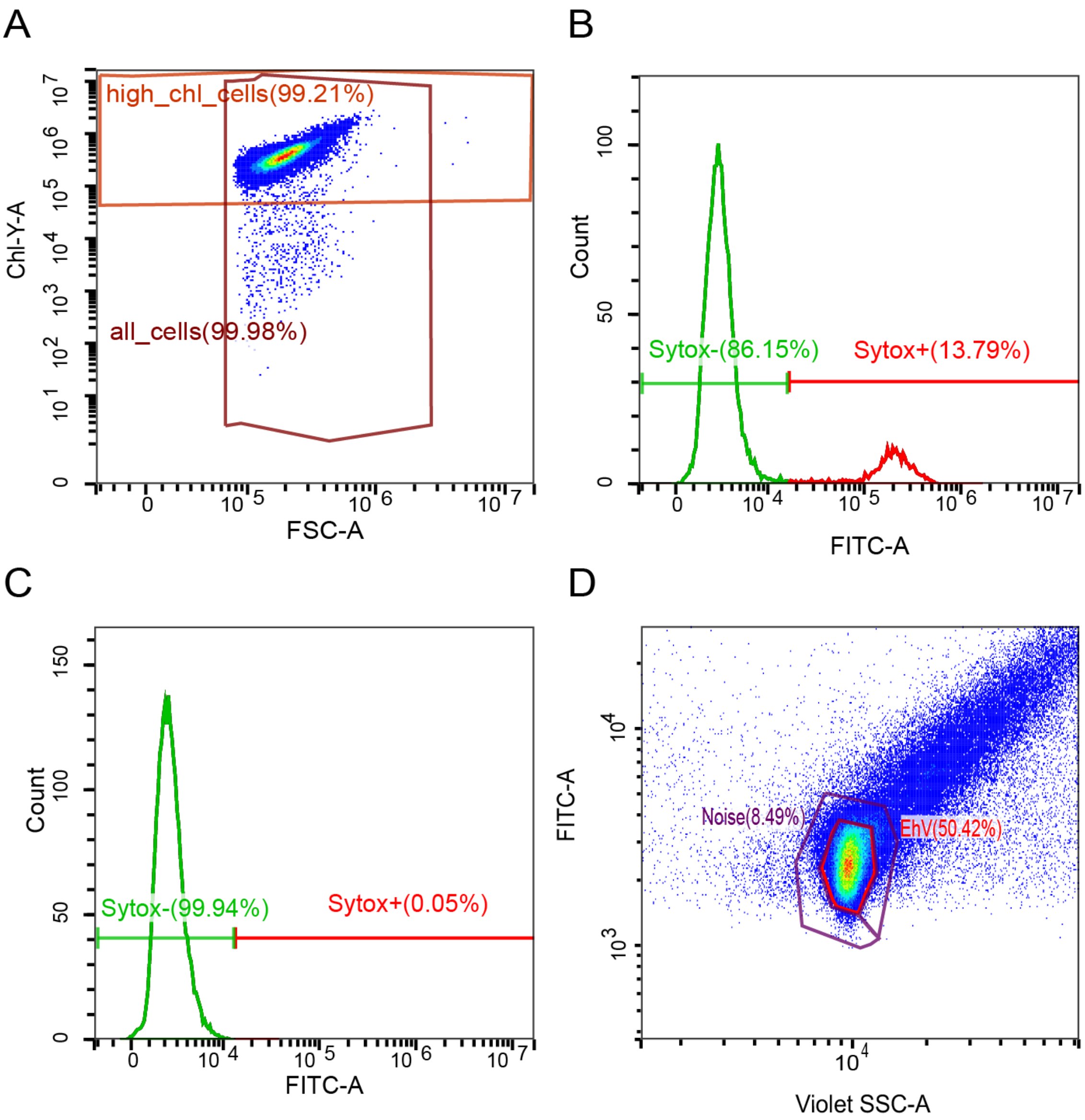


**Figure S14. Gating of algal cell, dead cell and virus particle populations by flow cytometry**. (A) Representative dot plot showing exponentially growing, non-infected algal cells. The quantification of live cells is based on the gate "high_chl_cells" (chl, chlorophyll). The y-axis represents chlorophyll autofluorescence and the x-axis represents forward scattered (FSC) light. (B-C) Histograms showing the percentage of dead cells ("Sytox+") in (B) a SYTOX-stained and (C) an unstained *E. huxleyi* 2090 culture infected with EhV-201 based on the gate "all-cells" in (A). The y-axis represents counted events (cells), and the x-axis represents green fluorescence by SYTOX staining (FITC). (D) Representative dot plot of viral particles stained with SYBR Gold. The inner gate confines extracellular viral particles. The outer gate measure background noise. The y-axis represents green fluorescence by the SYBR staining and the x-axis represents side scattered (SSC) light. Fluorescence is measured in arbitrary units.

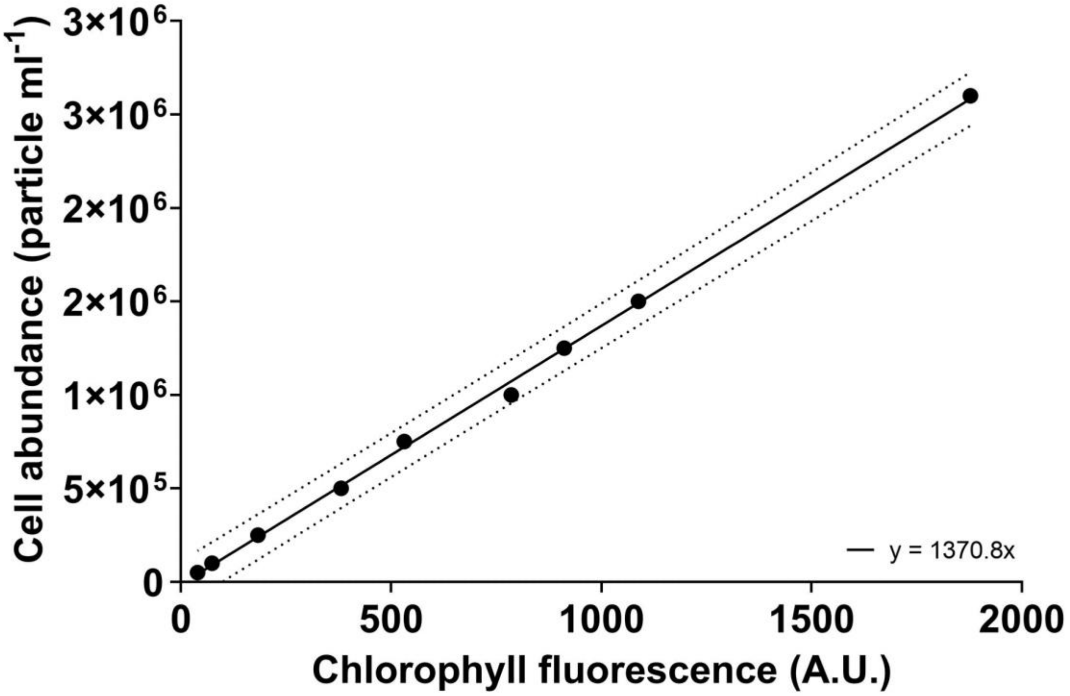


**Figure S15. Calibration curve to estimate cell abundances by measuring chlorophyll fluorescence using a plate reader**. The y-axis represents cell concentration as measured by flow cytometry. The x-axis represents the chlorophyll fluorescence emission (ex: 480 nm, em: 680 nm) of five measurements of the corresponding dilution. $R^2$ = 0.9994.

**Table S1. Percentages of cells that successfully generated monocultures following single-cell sorting.**

| Sorted strain or culture | Number of sorted single cells | Final size of monoculture collection | Percent of cells generating stable monocultures |
|---|---|---|---|
| 2090 | 485 | 123 | 25% |
| 2090-Rec | 388 | 74 | 19% |
| Rec-17 | 348 | 68 | 19% |
| Rec-32 | 348 | 43 | 12% |
| Rec-53 | 348 | 58 | 17% |
| 2090-BD5 | 288 | 38 | 13% |

Monocultures that did not survive monthly dilution as part of the culture maintenance were not included in the calculation.

**Table S2. Strains of *E. huxleyi* used to propagate the corresponding EhV strains.**

| EhV strain | *E. huxleyi* strain |
|---|---|
| EhV-201 | 2090 |
| EhV-86 | 374 |
| EhV-163 | 374 |
| EhV-ice01 | 374 |
| EhV-M1 | 374 |
| EhV-Rec | 2090-Rec |
